# Cross-domain competition for memory reactivation resources during sleep

**DOI:** 10.64898/2026.11.20.768209

**Authors:** Ainsley Temudo, Judith Nicolas, Bradley R. King, Genevieve Albouy

## Abstract

Targeted Memory Reactivation (TMR) enhances the consolidation of both declarative and procedural memories. As previous research highlights the importance of cross-domain interactions in shaping memory consolidation, we investigated here whether interacting memories from different domains can be co-reactivated by TMR during post-learning sleep, and in turn enhance consolidation. In a fully within-subject design, twenty-four young, healthy participants learned both a motor and an object sequence learning task that were linked or not by a shared ordinal structure. TMR cues associated only with the object task were presented during post-learning non-rapid eye movement (NREM) sleep which was monitored with electroencephalography (EEG) during a 90-minute nap. We tested whether reactivating object memories would indirectly reactivate the linked motor memory. Behavioral results showed no overall effect of association but indicated that TMR disrupted object memory consolidation without indirectly affecting performance on the linked motor task. Sleep EEG data point to a competition for reactivation resources whereby slow oscillation (SO; 0.5–2 Hz) and spindle activity (12–16 Hz) involved in both memory protection and potentiation during sleep were preferentially allocated to the covert consolidation of the motor trace at the expense of object memory. These findings suggest that cross-domain memories sharing a common learning structure compete for consolidation resources during post-learning sleep.

## Introduction

Optimizing learning and memory retention has been the focus of decades of research in neuroscience (see, e.g., Feld & Diekelmann, 2015; Phipps et al., 2021, for reviews). A well-studied experimental intervention that has shown promise to optimize consolidation in both the declarative and the procedural memory domains is Targeted Memory Reactivation (TMR). TMR is achieved by associating the learned information with sensory stimuli and replaying these stimuli offline, during the consolidation period, in order to reactivate the associated memory (Hu et al., 2020). Studies applying TMR during post-learning sleep in young adults have consistently demonstrated enhanced consolidation of both declarative and procedural memories (e.g., (Cairney et al., 2018; Cousins et al., 2016; Fuentemilla et al., 2013; Ngo & Staresina, 2022; Nicolas et al., 2022, 2025; Schreiner et al., 2024; Wang et al., 2019). At the neural level, electroencephalography (EEG) recordings show that TMR-induced memory reactivation is associated with modulations in sleep spindles (brief bursts of high frequency activity in the 12–16 Hz range; Cairney et al., 2018) and slow oscillations (SOs; slow, high amplitude waves in the 0.5–2 Hz range; Nicolas et al., 2022) characteristics, as well as increased coupling between these oscillations (Cairney et al., 2018; Nicolas et al., 2022).

While informative, this previous TMR research used tasks learned in isolation. Recent studies examining more complex learning processes tested whether multiple memories acquired simultaneously can be co-reactivated by TMR. Results suggested that TMR can simultaneously strengthen the consolidation of multiple memory traces from the same domain when they are linked through shared features during encoding, such as categorical relationships or co-occurrence (Schechtman et al., 2021, 2023). This process is supported by a modulation of sigma oscillations during reactivation such that greater TMR-induced sigma power predicts better memory retention (Antony et al., 2018; Schechtman et al., 2021). In addition, the co-reactivation of multiple memories during sleep results in power modulations in the delta–theta (1–8 Hz) and sigma (11–16 Hz) bands that scale to the number of associated items, suggesting that larger memory sets elicit greater oscillatory response amplitude (Schechtman et al., 2021). Altogether, these findings suggest that TMR can be used as a tool to reactivate multiple memories during post-learning sleep (Antony & Schechtman, 2024).

Despite the recent efforts reviewed above to examine multiple-memory processes, TMR research has largely continued to examine the declarative and procedural memory systems separately. This is surprising as real-world memories typically integrate components from these different domains (e.g. recalling a series of musical notes with the corresponding motor actions). Emerging theoretical frameworks in fact suggest that memory systems are less segregated than previously assumed and instead rely on overlapping neural networks and shared consolidation processes (Albouy et al., 2013; Brown & Robertson, 2007; Henke, 2010). Consistent with this view, recent work suggests that memories from different domains can interact during learning through, for example, a shared ordinal structure (Mosha & Robertson, 2016; Mutanen et al., 2020; Thong et al., 2026b, but see Thong et al., 2026a for a failed replication) and that these interactions can influence the fate of these memories. However, it remains unknown whether such cross-domain associations can be leveraged using TMR to modulate memory consolidation.

In the present study, that was pre-registered in the Open Science Framework (https://osf.io/), we used a TMR intervention to test whether memories from different domains - linked together via higher-order associations (i.e., a common ordinal structure between tasks) during encoding - would be reactivated together. In a within-subject design, twenty-four young healthy participants (13 females) were trained on both a motor sequence learning task and an object sequence learning task (procedural and declarative memories, respectively) that were – or were not – linked during encoding with higher-order associations (Figure 1, Temudo et al., 2026, 2025). TMR was administered during post-learning non-rapid eye movement (NREM) sleep with auditory cues corresponding to only one memory domain (Figure 2). We tested whether the memory from the other domain could be indirectly reactivated via cross-domain associations (Figure 2B). We hypothesized that TMR would enhance the consolidation of memories cued during sleep regardless of association. We also expected that this benefit would extend to memories in the other domain that were not directly cued during sleep but that were associated to the cued memory during learning via a common ordinal structure. We also sought to characterize the neurophysiological mechanisms supporting cross-domain memory reactivation. To this end, we analyzed electroencephalography (EEG) data acquired during TMR and examined modulation of specific electrophysiological markers of brain plasticity during sleep (spindles and SOs, and their coupling; Cairney et al., 2018; Nicolas et al., 2022). We predicted that these electrophysiological markers would be enhanced during the reactivation of multiple memories linked via higher-order associations, potentially reflecting co-reactivation processes.

**Figure 1.**
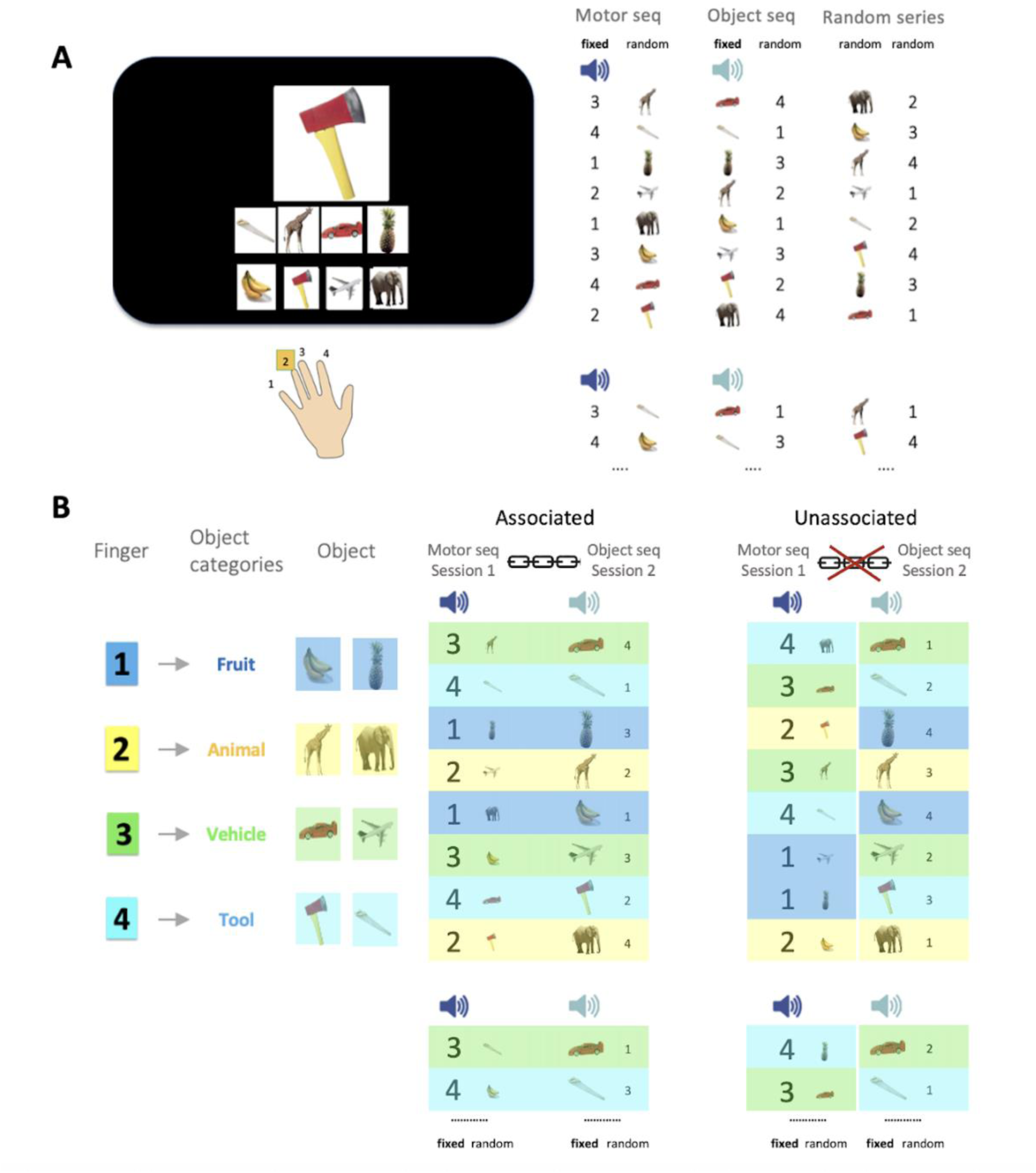
*Experimental task and conditions**. A.** Serial reaction time task (SRTT)* used in our earlier research to examine sequence learning in the two different memory domains (procedural vs. declarative) (Temudo et al., 2026, 2025). Left panel. A stimulus appears in the center of the screen and participants are instructed to respond as fast and as accurately as possible according to the key/object mapping displayed at the bottom of the screen. The key-object mapping changed after each stream of 8 elements (i.e., every 8 objects / 8 key presses) to orthogonalize motor and object sequence learning within each learning session (see right panel). Right panel. Three different task conditions were designed: a motor sequence (seq) with a fixed repetition of a specific pattern of key presses and random object presentation, an object sequence with fixed object presentation and random key presses and a random series with random key presses and object presentation (see methods). A distinct sound was played at the start of each sequence repetition for both the motor and object sequences. In both panels, numbers represent fingers, whereby 1 and 4 correspond to little finger and index finger of the participants’ non-dominant hand respectively. The object and motor sequences were learned in two separate sessions (see panel B and Figure 2A). **B.** *E*x*perimental conditions.* Left panel: Finger-object category mapping used to create higher-order associations between motor and object sequence tasks. Each finger in the motor sequence (1 to 4) is associated to a specific object-category (fruit, animal, vehicle, tool) in the object sequence. Right panel: In a within-subject design, participants completed both an associated (depicted by a chain link) and an unassociated visit (depicted by a crossed-out chain link). In the associated condition, object and motor sequences were linked between learning sessions through the fixed finger-object category mapping shown on the left panel (e.g., key 4 mapped to the tool category between learning sessions). In the unassociated condition, object and motor sequences were not linked between sessions, i.e. fingers were mapped to objects from different categories between sessions (e.g., key 4 mapped to vehicle and fruit categories between learning sessions). In both panels, object categories and fingers are indicated by different colors (blue = fruit/1, yellow = animal/2, green = vehicle/3, cyan = tool/4; see left panel) to highlight cross-domain associations (or the lack thereof) between learning sessions. Example pairs of object and motor sequences are shown for both visits, see supplemental Table S1 for all sequences and sequence pairs used in the experiment. All object images were derived from an open access online database at http://olivalab.mit.edu/MM/uniqueObjects.html (Brady et al. 2008). The picture of the hands is taken from an open access online sharing service at https://www.clker.com.

**Figure 2.**
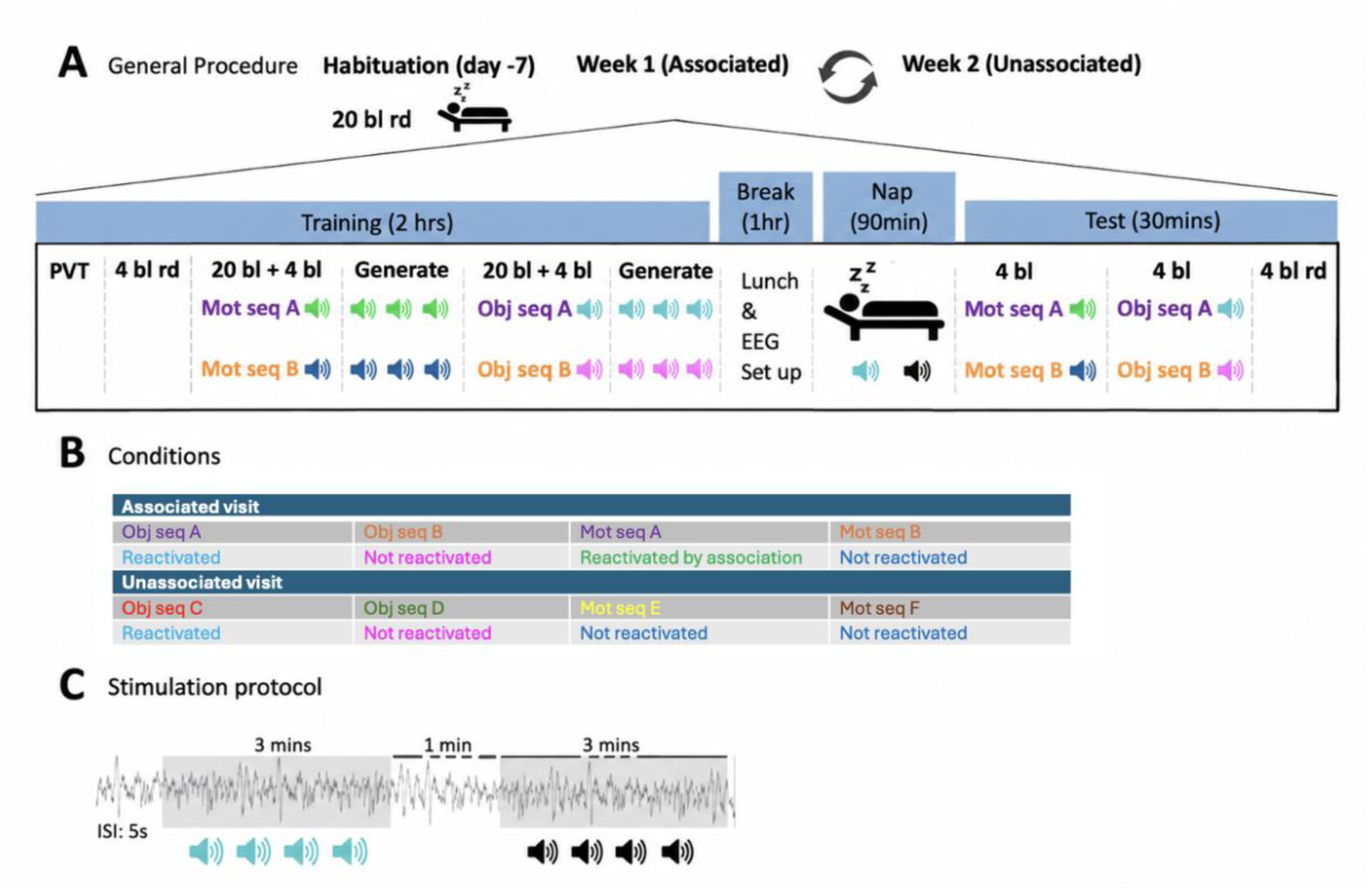
*Experimental design*. **A.** *General procedure*. After a habituation session including a habituation nap and practice of the random (rd) serial reaction time task (SRTT; ∼1 week prior to the experiment), participants completed 2 experimental visits (associated/unassociated, order counterbalanced across participants) ∼1 week apart. Each experimental visit started with an objective assessment of alertness using the Psychomotor Vigilance Task (PVT), followed by 4 blocks (bl.) of the random condition and the auditory threshold test (see methods). Participants were then trained on 2 motor sequences (20 blocks of training followed by 4 [pre-nap] test blocks for each motor sequence; see description of the conditions in panel B) each associated with a different auditory cue. Participants then performed a generation task in which they were asked to produce the learned sequences of movements when the corresponding auditory cues were presented to assess sequence and sequence-cue association knowledge. Next, participants were trained on 2 object sequences (20 blocks of training followed by 4 [pre-nap] test blocks for each object sequence; see description of the conditions in panel B) each associated with a different auditory cue. This was followed by a generation task similar to that described above. Participants’ brain activity was then monitored with EEG during a 90-minute nap opportunity during which one of the two object sequences was reactivated (*reactivated* object memory, see panel B) by replaying the corresponding auditory cue (*memory cue;* cyan). Sounds corresponding to the second object sequence were not played during the post-training nap and this condition therefore served as a *not reactivated* condition (pink; and see panel B). Sounds corresponding to the motor sequences (green and dark blue) were not presented during the nap. These conditions allow us to test whether a motor memory trace can be *reactivated by association* (or *not reactivated in the absence of association*) via higher-order associations with the other memory domain (see panel B). Another sound that was not associated with learning was also played and used as a *control cue* (black). After the nap, participants were tested on 4 [post-nap] retest blocks of each of the 4 sequences previously learned (always motor first followed by object second) followed by 4 blocks of the random condition. The figure depicts the design for the associated visit in which motor and object sequences are associated via higher-order associations (Motor sequence A is associated to Object sequence A and Motor sequence B to Object sequence B). The unassociated experimental visit followed the same design but with different sequences and sounds (e.g., Motor sequences C and D that are not associated with Object sequences E and F). **B.** *Visits and conditions.* Exemplar sequence combinations for a participant and the corresponding reactivation conditions (assignment of sequences to reactivation condition and visit were counterbalanced). The associated visit involved two reactivated conditions, e.g., *reactivated* [Obj seq A] and *reactivated by association* [Mot seq A] which corresponds to a memory reactivation load of two traces. In contrast, the unassociated visit involved only one reactivated condition, e.g., *reactivated* [Obj seq C] resulting in a memory reactivation load of one trace. **C.** *Stimulation protocol*. Sleep was recorded using an 11-channel EEG setup during the nap period (see methods). Sleep stages were monitored in real time, and auditory cues were delivered during NREM sleep stages 2 and 3. Stimuli were presented during three-minute stimulation intervals of each cue type (*memory* or *control*) alternating with a silent one-minute period (rest intervals). The inter-stimulus interval (ISI) was 5s. The stimulation was manually stopped when REM sleep, NREM1 or wakefulness was detected. Sleep metrics derived from offline scoring, along with the distribution of auditory cues in the different sleep stages for both sessions are reported and compared in Supplementary Tables S2–S3. The sleep data show that participants slept for a substantial portion of the recording period across both visits (average total sleep time: 69.4 min, mean sleep efficiency: 78.7%), spent on average 58.1 min in NREM2-3 sleep and auditory cues were delivered during NREM2-3 sleep with high accuracy (mean stimulation accuracy: 95.3%). None of these metrics differed between visits (see supplements).

## Results

### TMR disrupts the consolidation of the reactivated object memory but not the consolidation of the motor memory reactivated by association

We tested whether memories from different domains linked through higher-order associations during encoding showed greater TMR-induced consolidation than memories not linked via such associations. To do so, offline changes in performance speed were computed between the pre-nap test session and the post-nap retest session for both tasks (object vs. motor), reactivation (reactivated vs. not reactivated) and visit (associated vs. unassociated) conditions (see Figure 3A; note that in panel A, performance improvement is reflected by a decrease in response time RT). A significant task × reactivation condition interaction was observed (F(1,23) = 4.22, p = .05, ηp² = .16) but contrary to our expectations, results revealed that offline changes in performance speed tended to be greater for the non-reactivated sequence compared to the reactivated sequence in the object task (object, reactivated vs. non reactivated: F(1,23) = 3.48, p = .08, ηp² = .13) while no such effect was observed for the motor task (motor, reactivated vs. non reactivated: F(1,23) = 0.71, p = .41, ηp² = .03; Figure 3B, note that in panel B, a positive offline change in performance reflects overnight performance improvement). These effects did not interact with the visit factor (task × reactivation x visit: F(1,23) = 0.06, p = .81, ηp² = .003; see Supplemental Section 1B for a complete report of the statistical results on performance speed and accuracy and Figure S1B for a display of performance accuracy).

**Figure 3.**
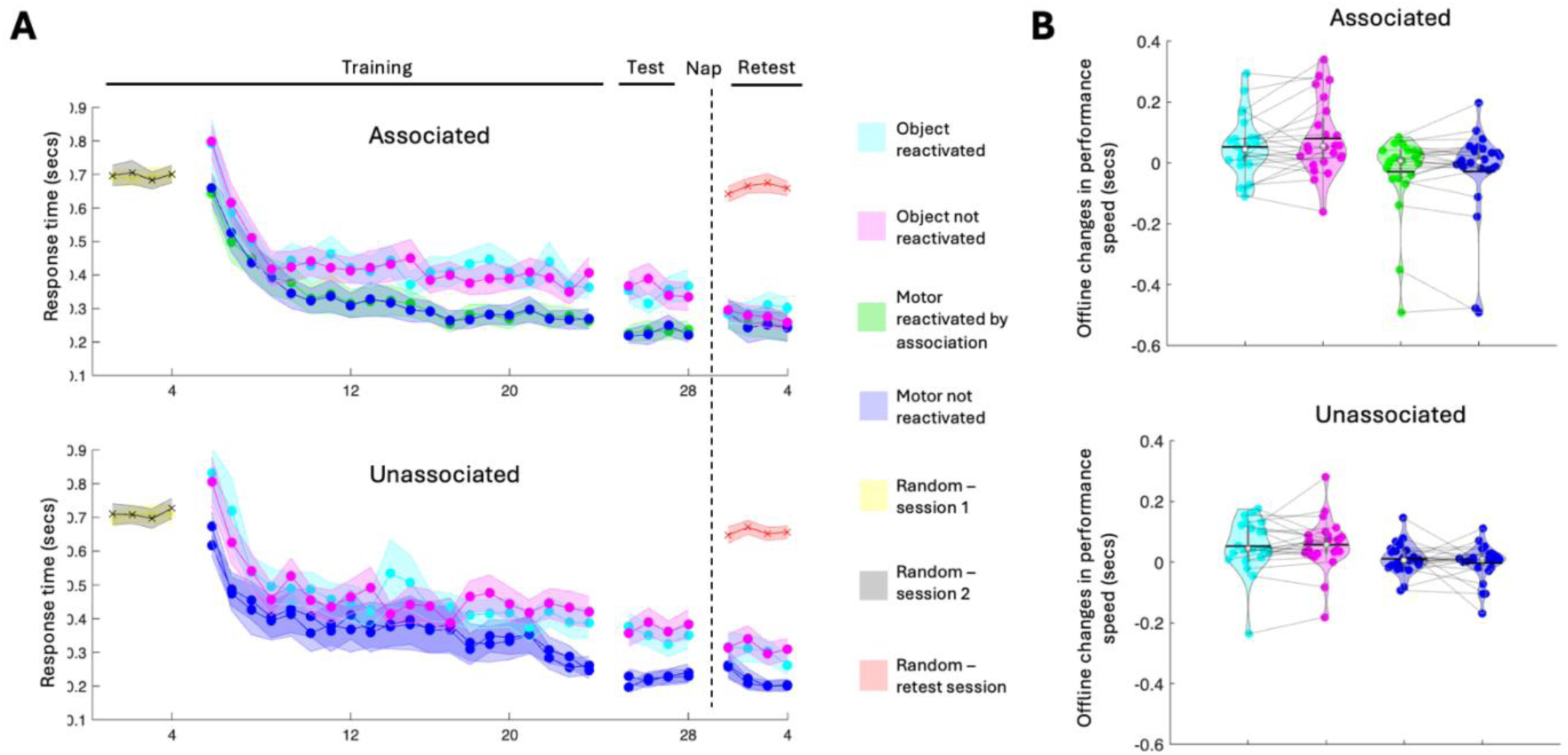
*Performance speed*. **A***. Performance speed during training.* Response time (RT, in seconds) across blocks for each sequence (reactivated/non-reactivated), each task (motor/object) and each visit (associated/unassociated). Performance speed did not differ between reactivated and non-reactivated sequences across tasks and visits prior to sleep (see Supplementary Sections 2-3 for a complete report of the statistical analyses related to performance prior to sleep). Crosses and circles indicate blocks of the random and sequence conditions, respectively. Shaded error bars=SEM, dotted line indicated when the nap occurred (N = 24). **B**. *Offine changes in performance speed* (in seconds) were calculated as the average response time across the 4 pre-nap test blocks minus the average across the 4 post-nap retest blocks, for both sequences (reactivated/non-reactivated) of each task (motor/object) and visit (associated/unassociated) conditions. Results indicate that TMR disrupted the consolidation of the reactivated object memory while no effect was observed on the consolidation of the motor memory reactivated by association (task × reactivation: F(1,23) = 4.22, p = .05, ηp² = .16; reactivated vs. non reactivated, object: F(1,23) = 3.48, p = .08, ηp² = .13; motor: F(1,23) = 0.71, p = .41, ηp² = .03). Colored circles represent individual data, jittered in arbitrary distances on the x-axis to increase perceptibility. Grey lines map individual datapoints across conditions. White circles represent medians, and black horizontal bars represent means. The shape of the violin depicts the kernel density estimate of the data (N = 24).

Taken together, these results suggest that higher-order associations did not influence memory consolidation. However, TMR disrupted the consolidation of the reactivated object sequence but did not modulate the consolidation of the motor sequence reactivated by association.

### Control cues elicited greater auditory-evoked responses and spindle activity under high memory load

EEG data collected during the TMR episode were analyzed to test whether memory (vs. control) cueing modulated auditory-evoked responses and the characteristics of electrophysiological markers critically involved in memory reactivation during sleep, i.e., SO and spindle events, as well as sigma oscillations, taken as another proxy of spindle activity (Nicolas et al., 2022, 2025; Cairney et al., 2018). We further examined whether these effects differed between associated and unassociated visits (i.e., whether effects varied as a function of memory reactivation load).

Auditory-cue locked analyses were performed on each EEG channel to examine the effects of auditory stimulation conditions (memory vs. control cue) on both event-related potentials (ERP) and oscillatory activity (time–frequency analyses), and whether these effects were modulated by higher-order associations (associated vs. unassociated; see Supplemental Table S3 and methods for information about the number of auditory events for each cue and visit conditions). Figure 4A shows ERPs for each cue and visit conditions. Cluster-based permutation (CBP) tests (Maris and Oostenveld 2007) on the memory vs. control cue comparison did not reveal any significant spatio-temporal ERP clusters in either visit (Figure 4B-C). In contrast, a significant interaction cluster ([memory – control] x [associated – unassociated]) was observed approximately 1s post-cue onset whereby ERP amplitude was greater for the control compared to the memory cue in the associated visit while the opposite pattern was observed in the unassociated visit (Figure 4D; *cluster a* - p = .01, Cohen’s d = -.69, 0.8 -1s relative to cue onset, cluster spanning all channels excluding FPz, F4, P4 and Oz). CBP tests conducted on the time-frequency representations (TFR) across time, frequency, and channels did not reveal any significant clusters either within visits or when comparing the associated and unassociated visits (see TFRs of power in Figure 4B-D and Supplementary Figure S2 for TFRs plotted for each cue and visit conditions). Together, these results suggest that the control sounds that did not carry memory information elicited stronger auditory-evoked responses under high memory load, i.e., during the associated visit when two associated memory traces were potentially reactivated during sleep.

**Figure 4.**
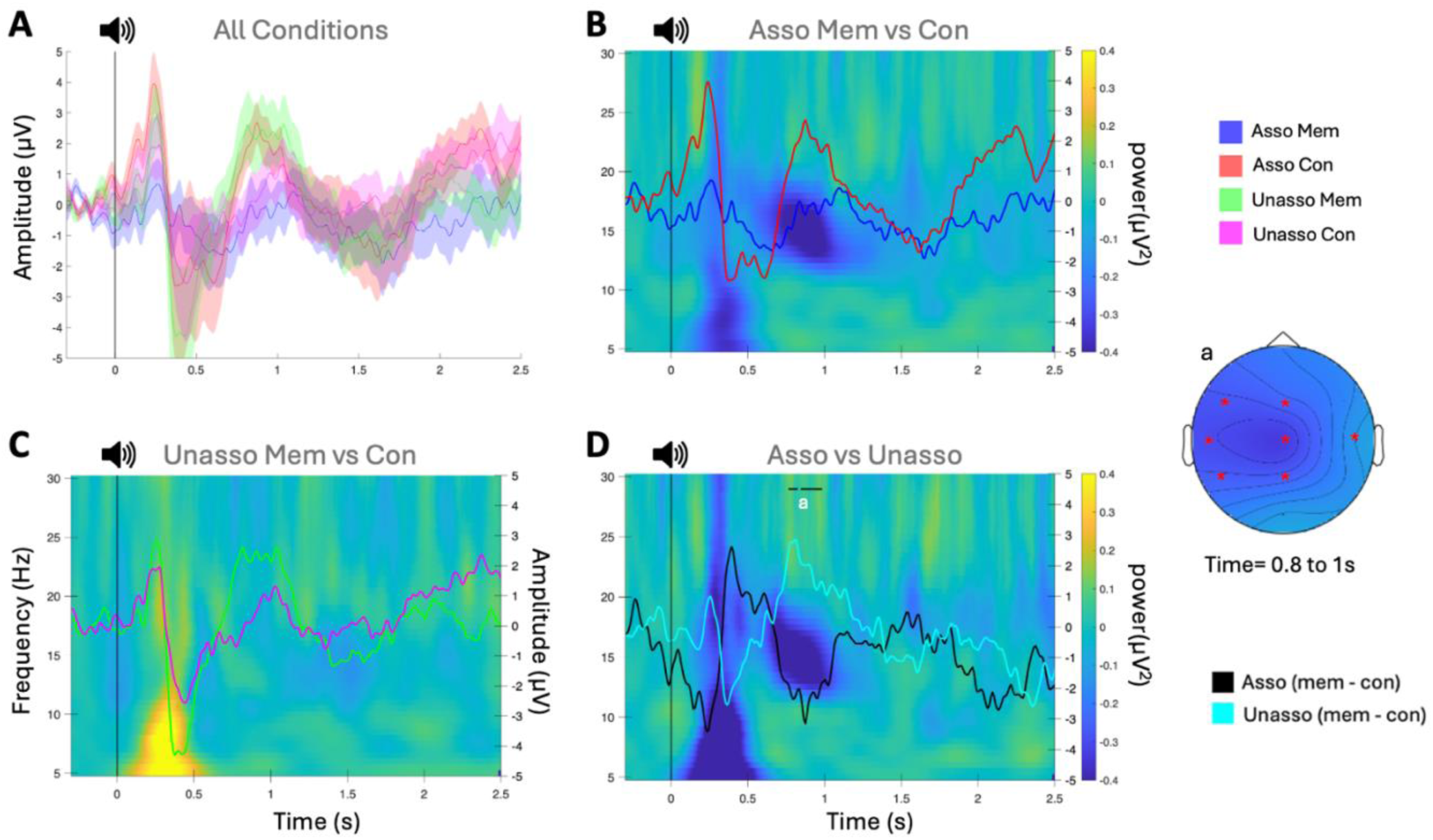
*Auditory cue-evoked responses during sleep within and between associated and unassociated visits.* Event-related potentials (ERPs; line plots displayed for channel Cz) are overlaid on time–frequency representations (TFRs; 5–30 Hz) of EEG power (displayed for channel Cz), time-locked to auditory cue onset (0 s; black vertical line)., Asso: associated, Unasso: unassociated, Mem: memory, Con: control. **A.** ERPs are shown separately for associated memory (blue), associated control (red), unassociated memory (green), and unassociated control (pink) cues. Shaded regions represent SEM. **B.** Difference between memory and control cues in the associated visit (N=24). No significant clusters were observed for either ERP or TFR analyses. **C**. Difference between memory and control cues in the unassociated visit (N=23^¥^). No significant clusters were observed for either ERP or TFR. **D**. Difference between the associated and unassociated (N=23^¥^) contrasts (associated memory – control [black]) − (unassociated memory – control [cyan]). A significant spatio-temporal cluster (a) was observed for ERP at 0.8–1.0 s post-cue (p = .01; Cohen’s d = -.69) across all channels excluding FPz, F4, P4 and Oz. No significant clusters were observed for TFR. Black horizontal lines indicate the time points belonging to the significant spatiotemporal clusters in ERP amplitude difference identified through cluster-based permutation tests. Topographical map displays the spatial clusters highlighted by CBP, with red stars marking channels included in the significant cluster. ^¥^ Data is missing for one participant in the unassociated visit due to experimental error resulting in stimulation triggers not being recorded for this participant.

SO-trough locked analyses (both ERP and TRFs) were then conducted to examine whether the different stimulation conditions (as compared to non-stimulation [rest] intervals) modulated SO-related neural responses differently between the associated and unassociated visits (see Supplemental Table S4 for number of SO events in each interval and visit; and Supplementary Figure S3 for a depiction of the responses on the separate conditions, i.e., associated memory cue, associated control cue, associated rest, unassociated memory cue, unassociated control cue, unassociated rest). CBP tests were first applied in a pairwise fashion to identify differences in SO amplitude between conditions within each visit. In the associated visit, SO amplitude did not differ between the memory vs. control cue stimulation intervals. However, CBP analyses showed that, during the associated visit, SO amplitude at the trough of the oscillation was greater during periods of no stimulation compared to periods of both memory cue stimulation (Figure 5A; *cluster a*: −0.1 to 0.2 s, *p* = .02, Cohen’s d= .86, FPz cluster) and control cue stimulation (Figure 5A; *cluster b*: -0.1 to 0.2 s, *p* = .006, Cohen’s d = .86; *cluster c*: 0.02 to 0.2 s, *p* = .006, Cohen’s d = .95, frontal clusters). No significant spatio-temporal clusters were observed for the unassociated visit or when comparing ERPs between visits (Figure 5B). Overall, these results suggest that auditory stimulation - irrespective of the cue condition - decreased SO amplitude under high memory load, i.e., during the visit when the associated memory traces are potentially reactivated together. Note that this effect was not different between visits and that exploratory analyses indicated that this effect was significant across association conditions (Supplementary Figure S4).

**Figure 5.**
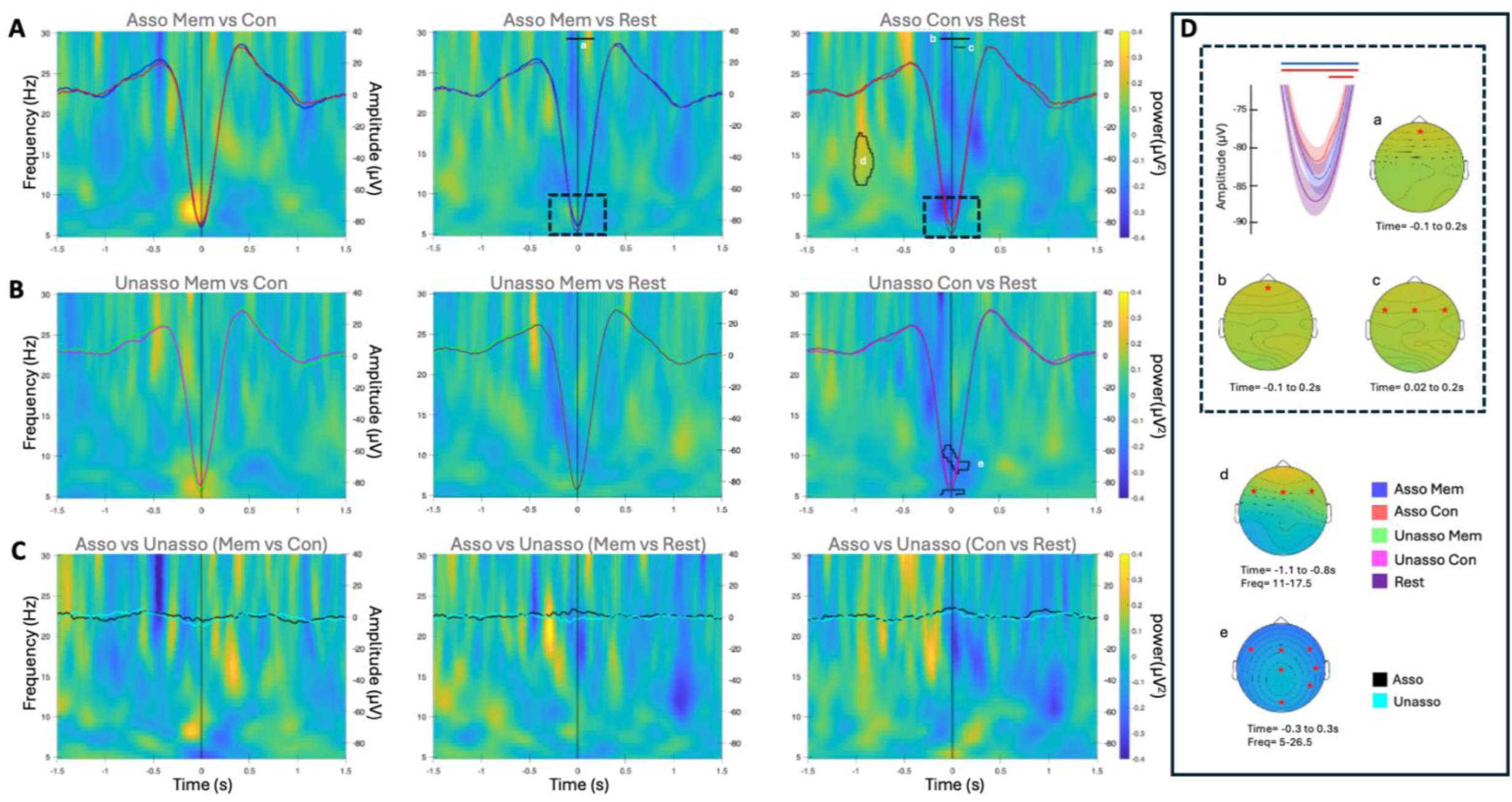
*SO-locked responses during sleep within and between associated and unassociated visits*. Event-related potentials (ERPs; line plots displayed for Fz channel) overlaid on time–frequency representations (TFRs; 5–30 Hz) of EEG power (displayed for channel Fz), time-locked to SO trough (0 s; black vertical line). Asso: associated, Unasso: unassociated, Mem: memory, Con: control. **A.** Associated visit (N=24); *left panel*; difference between memory (blue) and control (red) cue stimulation; *middle panel*; difference between memory cue stimulation and rest (purple) showing a significant spatiotemporal cluster (a) from −0.1 to 0.2 s (p = .02, Cohen’s d = .86) indicating greater SO amplitude during rest compared to memory cue stimulation; *right panel*; difference between control cue stimulation and rest showing significant spatiotemporal clusters (b) from -0.1 to 0.20 s (p= .006, Cohen’s d = .86) and (c) from 0.02 to 0.2 s (p = .006, Cohen’s d = .95) indicating greater SO amplitude for rest compared to control cue stimulation intervals. TF analyses revealed a significant spatiotemporal-frequency cluster (d) from −1.1 to −0.8 s, 11–17.5 Hz (p = .01, Cohen’s d= .96) showing greater sigma power 1s before the SO-trough during control stimulation as compared to rest. See panel D for a description of the SO amplitude effects and for an inset zooming on SO trough and the frontal topography of the reported ERP and TF effects. **B.** Unassociated visit (N=23^¥^); *left panel*; difference between memory (green) and control (pink) cue stimulation; *middle panel*; difference between memory cue stimulation and rest (purple); *right panel*; difference between control cue stimulation and rest. No differences in ERP were observed but TF analyses revealed a significant spatiotemporal-frequency cluster (e) from −0.3 to 0.3 s, 5–26.5 Hz (*p* = .004, Cohen’s d= .-0.39) indicating lower power for control cues compared to rest (see panel D for topography of this effect). **C.** Comparison between associated (N=23^¥^; black) and unassociated (N=23^¥^; cyan) visits didn’t yield any significant results; *left panel*; memory vs. control cue stimulation; *middle panel*; memory cue stimulation vs. rest and *right panel*; control cue stimulation vs. rest. Black horizontal lines indicate time points belonging to significant spatiotemporal clusters in SO amplitude difference identified through cluster-based permutation tests. Black contours in the TFR maps highlight significant spatial time–frequency clusters. **D.** Inset showing zoom on SO trough in the associated visit where the rest condition [purple] showed greater SO amplitude compared to the two auditory stimulation conditions (memory [blue] and control [red]; shaded regions represent SEM). Topographical maps display the spatial distribution of all CBP effects, with red stars marking channels included in each significant cluster. ^¥^ Data is missing for one participant in the unassociated visit due to experimental error resulting in stimulation triggers not being recorded for this participant.

With respect to SO-trough locked oscillatory power, CBP tests on the associated visit revealed that sigma power was greater during periods of control stimulation compared to periods of no stimulation approximately 1 s prior to the SO trough (Figure 5A; *cluster d*: −1.1 to −0.8 s; 11–17 Hz; *p* = .01, Cohen’s d= .96, cluster spanning frontal channels). In the unassociated visit, a significant negative cluster was observed for the control vs. rest contrast around the SO trough (Figure 5B; *cluster e*: −0.3 to 0.3 s; 5–26.5 Hz; *p* = .004, Cohen’s d = -0.39, cluster spanning Fz, Cz, Oz, F3, F4, C4 and P4) reflecting decreased theta-alpha-band power for control cues compared to rest. No significant clusters were identified when comparing oscillatory activity between visits. In line with the auditory-ERP results, these findings therefore indicate that control cues elicited greater sigma power as compared to rest before the trough of the SO. Note that these effects did not survive comparisons between visits.

Last, SO and sleep spindle events were automatically identified from the stimulated and not-stimulated intervals using an established detection algorithm (Vallat & Walker, 2021) used in our previous research (e.g., Nicolas et al., 2022, 2025) and corresponding results are reported in the Supplementary Section 4 and Supplementary Figure S5. In summary, the data indicate that auditory stimulation increased *SO density* irrespective of the auditory cue and visit conditions. In line with the sigma power results described above, spindle analyses showed that control sounds that did not carry memory information induced an overall increase in *spindle amplitude* that was greater during the associated vs. the unassociated visit, i.e. under high memory load.

Altogether, these electrophysiological findings suggest that, under higher memory load, i.e., during the associated visit when memory traces are linked via higher-order associations and potentially reactivated together, auditory stimulation resulted in decreased SO amplitude and control cues elicited greater auditory-evoked responses and enhanced spindle activity.

### TMR-induced modulations of SO amplitude and sigma power are related to a greater beneficial effect of reactivation by association on motor performance

Brain-behavior correlation analyses were conducted as per our pre-registration to assess the relationship between the effect of TMR on performance and EEG-derived measures. Specifically, for both the motor and object tasks, within each visit, we examined correlations between the TMR index (computed as the difference in offline changes in performance between the reactivated and not reactivated sequences) and the difference between memory and control cues for both auditory cue– and SO-locked responses (ERP and power). Cluster-based permutation (CBP) tests were applied to identify significant brain–behavior relationships.

No significant clusters were observed when correlating the TMR index with auditory cue–locked ERP or TFR. In contrast, significant clusters emerged for SO trough–locked analyses, specifically in relation to the motor TMR index in the associated visit. For SO amplitude, CBP analyses revealed a significant negative spatiotemporal cluster 1 s post SO-trough (black horizontal line in the middle panel of Figure 6) whereby greater SO amplitude difference between the memory and the control cue stimulation intervals (i.e., a more negative trough for the memory condition) was related to greater motor TMR index (Figure 6; *cluster a1:* from 0.68 to 1.01 s relative to the SO trough; *p* = .02, rho = -0.43, cluster spanning channels Cz, Fz, Pz, C3, P3 and P4). Exploratory analyses testing for a difference in brain-behavior correlations between the motor and the object TMR index revealed that the relationship described above was greater for the motor than for the object TMR index (*cluster a2:* from 0.75 to 1.03 s relative to the SO trough; *p* = .04, rho = -0.55, cluster spanning channels Cz, Fz, Pz and C4, [red horizontal line in the middle panel of Figure 6 and scatter plot related to cluster a]). Importantly, no such relationship - or difference in relationship - was observed during the unassociated visit.

**Figure 6.**
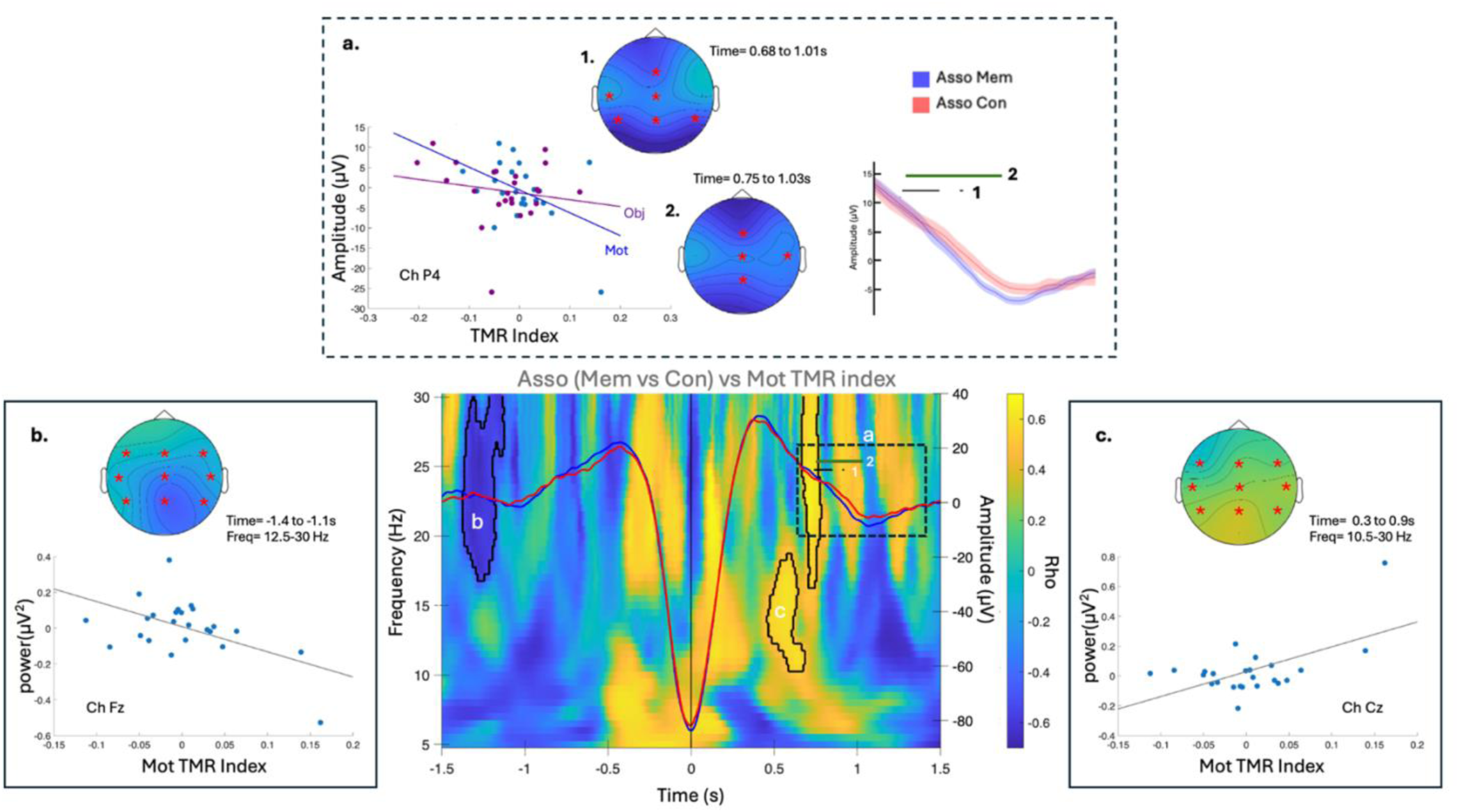
*Correlations between TMR index and SO trough-locked neural responses in the associated visit*. Asso: associated, Mem: memory, Con: control, Mot: motor, Obj: object, Ch: channel, Freq: frequency. Middle panel: SO grand averages (displayed for channel Fz) for the memory (blue) and control (red) cue stimulation intervals during the associated visit super-imposed on the time–frequency representation (time-locked to SO trough-0 s; black vertical line) of the correlation coefficients (rho) for the relationship between the motor TMR index (i.e., motor reactivated by association – motor not reactivated) and the oscillatory power difference (memory − control contrast) for the associated visit (N=24; displayed for channel Cz). The black horizontal line [a1] indicates time points belonging to the significant spatiotemporal cluster identified using cluster-based permutation tests showing a correlation between the difference in SO amplitude and the motor TMR index. The green horizontal line [a2] indicates time points belonging to the significant spatiotemporal cluster from the exploratory CBP analyses testing for a difference in correlation between the motor and object TMR indices. Inset a (dashed box) highlights time points belonging to significant spatiotemporal clusters of correlations, their topography and corresponding scatter plots (see description of the effects below). In the TFR, black contours indicate significant spatiotemporal-frequency clusters of correlations identified using cluster-based permutation tests. Clusters b and c indicate that sigma-beta power modulation between the memory and the control cue are related to a greater motor TMR index. Insets b and c (solid boxes) show the topography of the significant spatiotemporal-frequency clusters of correlations and the corresponding scatter plots (see description of the effects below). **Inset a**. Significant cluster a1 indicates that greater motor TMR benefit (blue) was associated with greater negative SO peak for memory cue compared to control cue (from 0.68 to 1.01 s; p= .02; rho = -0.43; scatter plot displayed for channel P4). Significant cluster a2 indicates that the correlation between SO amplitude difference and motor TMR index was greater than with the object TMR index (purple) (from 0.75 to 1.03 s; p = .04; rho = -0.55). Top panel displays a zoomed view of memory and control stimulation SO amplitude across the significant time window. **Inset b**. Cluster b indicates that greater power for the control cue compared to the memory cue 1 s before the SO-trough was associated with greater TMR benefit on the motor task (from −1.4 to −1.1 s in the 12.5–30 Hz range; p = .04; rho = - 0.58; cluster spanning all channels excluding FP and Oz; scatter plot displayed for channel Fz). **Inset c**. Cluster c indicates that greater sigma-beta power for the memory cue compared to the control cue at the SO-peak was related to a greater motor TMR index (from 0.3 to 0.9 s in the 10.5–30 Hz range; p = .02; rho = 0.57; cluster spanning all channels excluding Fz and Oz; scatter plot displayed for channel Cz). Topographical maps display the spatial distribution of CBP effects, with red stars marking channels included in each significant cluster. Note that the clusters highlighted above are no longer significant when removing the participant presenting the largest motor TMR index.

For oscillatory power, CBP analyses revealed two significant clusters with the motor TMR index within the associated visit. A significant negative cluster was identified in the sigma-beta band more than 1 s prior to the SO trough (Figure 6; *cluster b*: −1.4 to −1.1 s; 12.5–30 Hz; *p* = .04, rho = -0.58, cluster spanning all channels excluding FPz and Oz) and a significant positive cluster was observed in the sigma-beta band at the SO peak (Figure 6; *cluster c*: 0.29 to 0.85 s; 10.5–30 Hz; *p* = .02, rho = 0.57, cluster spanning all channels excluding FPz and Oz). Results for the negative cluster (b) indicated that greater power for the control cue compared to the memory cue 1 s pre-SO-trough was correlated with greater motor TMR index. In contrast, the positive cluster (c) showed that greater power for the memory cue compared to the control cue at the SO peak was associated with greater motor TMR index. No significant clusters were observed for the correlations with the object TMR index or during the unassociated visit.

Together, these results suggest that memory vs. control cue modulation of SO amplitude and sigma power at different phases of the SO are related to a greater beneficial effect of TMR by association on the consolidation of the motor memory trace.

## Discussion

This study tested whether reactivating a memory trace from one domain during sleep could indirectly benefit a linked memory from a different domain through higher-order associations acquired during learning. Contrary to our hypotheses, TMR did not produce the predicted cross-domain benefits such that reactivating the object memory did not influence the consolidation of the motor memory trace reactivated by association. Instead TMR tended to disrupt the consolidation of the cued object memory compared to the uncued object memory irrespective of association. Sleep EEG analyses revealed that auditory stimulation reduced SO amplitude which possibly contributed to the disruptive effect of TMR on object memory consolidation. EEG data also consistently showed heightened electrophysiological responses (auditory-locked ERPs, SO-locked sigma power and spindle characteristics) to the control cue in the visit with the higher memory load (i.e., when two memory traces linked via higher-order associations are potentially reactivated). Brain-behavior analyses revealed that SO and spindle activity was associated with the consolidation of the motor memory trace reactivated by association in two distinct ways that might reflect different processes. On the one hand, greater TMR-induced modulation of SO amplitude and sigma power at the peak of the SO were correlated with improvement in performance on the motor task, suggesting a role of SO and SO-peak nested sigma oscillations for the reinstatement of the motor memory trace by association. On the other hand, results showed greater spindle amplitude and SO-trough nested sigma power during control stimulation in the associated visit which might reflect a gating mechanism through which memory consolidation is protected against irrelevant information processing under higher memory load. Our data shows that individuals who exhibited stronger electrophysiological indices of both the reinstatement and protective gating processes demonstrated greater (covert) motor memory consolidation.

TMR tended to disrupt the consolidation of the directly reactivated object memory task irrespective of cross-domain associations. These results stand in contrast with prior work showing that TMR applied during post-learning NREM sleep consistently strengthens the consolidation of declarative memories (Hu et al., 2020). Unlike previous studies showing TMR-induced increases in SO characteristics (Nicolas et al., 2022, 2025), the current EEG data showed that auditory stimulation significantly decreased SO amplitude as compared to rest. We propose that such decrease in SO amplitude might have impaired the consolidation of the directly reactivated (as compared to not-reactivated) object memory trace. It remains unclear why our paradigm induced such a disruptive TMR effect on both performance and SO features. It is possible that the multi-task paradigm used in the current study to maximize interaction (and potentially association) between memory domains (Brown & Robertson, 2007; Kim, 2020; Mosha & Robertson, 2016; Temudo et al., 2026) resulted in competitive, rather than cooperative, cross-domain interactions that potentially interfered with object sequence learning. Previous work indeed suggests that the short time interval between tasks (<2h, Brown & Robertson, 2007) and the specific task order (motor followed by object learning, Temudo et al., 2026) used in the current research may have resulted in a weaker encoding of the object memory trace. Given prior work showing that, when multiple memories compete (e.g., interfere) during learning, TMR can impair rather than strengthen the consolidation of the weaker memories (Joensen et al., 2022; Oyarzún et al., 2017), our results are therefore in line with this competitive framework. Interestingly, previous studies have attributed such disruption to a competition for limited consolidation resources (Brown & Robertson, 2007; Joensen et al., 2022). According to this view, one might expect a trade-off whereby object memory disruption would be parallelled by (or even correlated with) improvement in performance on the motor task. To test this possibility, we performed exploratory correlation analyses across visits (as this competition framework does not appear to be modulated by cross-domain associations). While the correlation analyses performed on the behavioral outcomes did not support such view (see Supplementary Section 5 for analyses showing no relationship between motor and object changes in performance after TMR), the exploratory brain-behavior correlation analyses presented in Figure 7 showed a pattern of results compatible with the *competition for consolidation resources* account (Diekelmann et al., 2011; Rasch & Born, 2007). Specifically, the difference in SO amplitude between the memory and control cues across visits was differently related to motor and object TMR indices such that a greater amplitude difference was correlated with greater motor TMR index and lower object TMR index (Figure 7). Consistent with the concept of resource re-allocation during memory reactivation (Guttesen et al., 2026), these results therefore suggest that the greater the TMR-induced modulations of SO amplitude, the greater the TMR benefit on the motor task at the expense of the object task. Based on these observations, it is therefore tempting to speculate that a single memory cue can lead to drastically opposite effects mediated by SO amplitude modulation, i.e. the erasure of weak memories and the indirect potentiation of linked - and presumably more robust – memories from another domain.

**Figure 7.**
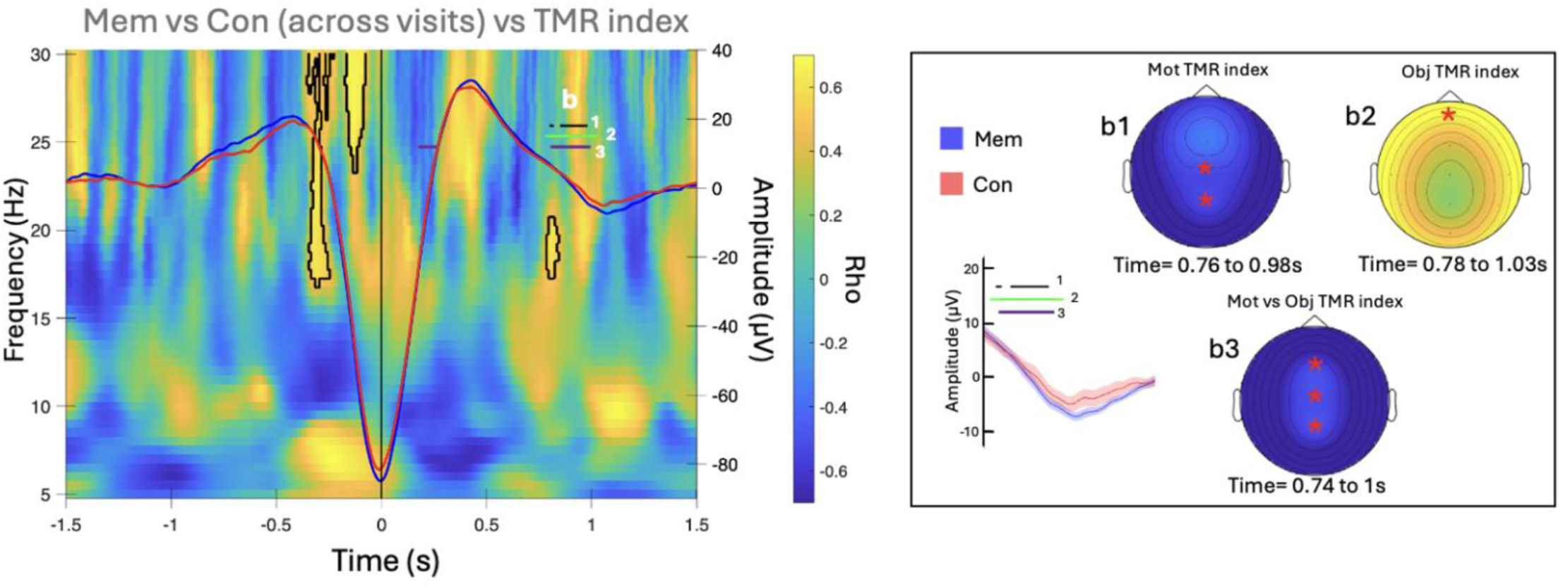
*Across visit exploratory analyses.* Mem: memory, Con: control, Mot: motor, Obj: object. Left panel. SO grand averages for the memory (blue) and control (red) stimulation intervals across visits super-imposed on the time–frequency representation of the correlation coefficients (rho) for the relationship between the motor TMR index (i.e., motor indirectly reactivated – motor not indirectly reactivated) vs. object TMR index (i.e., object directly reactivated – object not indirectly reactivated) and the oscillatory power difference (memory − control contrast) across visits (N=23**^¥^**; displayed for channel Fz). Horizontal line indicates time points belonging to the significant spatiotemporal clusters identified through cluster-based permutation tests. Cluster b1 (black line) indicates that greater motor TMR benefit is related to greater SO amplitude difference (b1: from .76 to .98 s; p = .3, spanning channels Cz & Pz), while Cluster b2 (green line) shows the opposite direction with object TMR benefit (b2: from .78 to 1.03 s; p = .02, channel FPz). Cluster b3 and (purple line) indicates that the correlation between the SO amplitude difference and the motor TMR benefit was greater than with the object TMR benefit (b3: from .74 to .1 s; p = .012, spanning channels Fz, Cz and Pz). TF analyses revealed significant spatiotemporal-frequency clusters (black contours) indicating greater power difference (memory – control) was correlated with greater TMR benefit difference (motor – object). Right panel: shows zoomed in SO amplitude highlighting significant spatiotemporal clusters of correlations (b clusters). Topographical maps display the spatial distribution of CBP effects, with red stars marking channels included in each significant cluster**. ^¥^** *Data is missing for one participant in the unassociated visit* due to experimental error resulting in stimulation triggers not being recorded for this participant.

While the effects discussed above were not modulated by higher-order associations between memory domains, our sleep EEG data unexpectedly indicated that responses to the *control cues* differed as a function of association. Specifically, auditory ERP amplitude was larger for the control cue as compared to the memory cue during the associated visit while an opposite trend was observed during the unassociated visit which resulted in a significant cue type by visit interaction. While the results of the associated visit are unexpected, the pattern observed in the unassociated visit is in line with previous TMR research showing larger potentials for cues carrying memory content compared to control cues (Cairney et al., 2018; Nicolas et al., 2022). Similar results were observed with markers of spindle activity. Specifically, spindle event amplitude was greater during control cue stimulation as compared to memory cue stimulation or rest intervals during the associated - compared to the unassociated - visit. Likewise, sigma power at the recurring through of the SO was greater in the control cue interval as compared to rest interval during the associated visit. These results are in line with our previous TMR research showing greater SO-trough nested sigma power for control as compared to memory cues during motor memory reactivation (Nicolas et al., 2022). Altogether, these results point to heightened electrophysiological responses to the control cue in the visit associated with the higher memory load (i.e., when memory traces are linked and potentially reactivated together). We suggest that these responses reflect a gating mechanism that protects memory consolidation against the processing of irrelevant stimuli (i.e., control sound not linked to any memory content) under a higher memory load. Sigma oscillations have indeed been proposed to shield ongoing consolidation from external interference and might reflect enhanced engagement of thalamocortical circuits involved in sensory filtering (Nicolas et al., 2022; Bandarabadi et al., 2020) when the memory load is greater. Interestingly, while these markers did not relate to performance on the object memory task - indicating that this protective mechanism did not preserve the directly reactivated (potentially weaker) object trace – they were correlated with performance on the indirectly reactivated memory trace (see below).

Brain-behavior correlation analyses indeed highlighted two different, presumably complementary, processes supporting the consolidation of the indirectly reactivated motor memory trace. First, as mentioned above, we observed that sigma bursts induced by control cues relative to memory cues at the trough of recurring oscillations were associated with greater TMR benefit on the motor task. Notably, no such relationship was observed with the object task or in the unassociated visit. These results are in line with our previous work showing a positive relationship between the power of control-cue-evoked sigma oscillations and the beneficial effect of TMR on motor memory consolidation (Nicolas et al. 2022). The current findings suggest that the sigma-related protective mechanisms described earlier - that are particularly heightened under higher memory load - may selectively shield the motor traces reactivated by association from interference induced by irrelevant stimulation. Second, our data show that greater sigma power at the peak of the SO for memory relative to control cues intervals predicted larger TMR-related improvements in motor performance. This also concurs with our previous research showing a positive relationship between TMR-induced SO-peak nested sigma oscillations and the beneficial effect of TMR on motor memory consolidation (Nicolas et al. 2022). We also observed here a similar correlation between the TMR-induced modulation of SO amplitude and the motor TMR index, and this relationship was significantly greater than with the object TMR index. Altogether, our results suggest that these established markers of memory protection and potentiation during sleep (Nicolas et al., 2022) were preferentially allocated to the covert consolidation of a motor memory trace linked via higher-order associations to the directly reactivated memory. It is worth noting that the brain-behavior correlation results discussed above should be interpreted with caution as they were no longer significant when removing the participants showing the largest motor TMR index.

In conclusion, the current data does not provide evidence for the co-reactivation of interacting memories during sleep. Instead, they suggest that memories from different domains sharing a common learning structure compete for resources supporting both memory reinstatement and protection against irrelevant information during sleep; two processes that critically contribute to memory consolidation. Future studies should systematically manipulate factors affecting encoding strength to determine the boundary conditions promoting cooperative - instead of competitive - consolidation that would enable cross-domain memories to be strengthened together during sleep.

## Material and Methods

This study was pre-registered in the Open Science Framework (https://osf.io/). Our pre-registration document outlined our hypotheses and intended analysis plan as well as the statistical models used to test our a priori hypotheses (available at https://osf.io/Agknw/). Deviations from the pre-registration are marked with a (#) and justifications for the deviations are presented in the Supplementary Section 6.

### Participants

Sample size computation was based on a previous TMR study from our group (Nicolas et al., 2022) showing a main effect of condition (reactivated vs. control; η² = 0.15) on offline gains in performance speed (motor task). The correlation coefficient was set to 0.5 (instead of 0.66 in the original study) to account for the different nature of the design (i.e., two tasks instead of one [motor & object] and 2 visits [associated & unassociated] as additional within-subject factors of interest). The primary contrast of interest was a *reactivation* (reactivated vs. not-reactivated, within-subject factor) by *visit* (associated vs. unassociated, within-subject factor) interaction using a repeated-measures ANOVA. The required sample size was 22 participants total (effect size f = 0.42, alpha = 0.05, power = 0.95). To ensure complete counterbalancing of the combination of sequences and sounds across participants (see details in the procedure section), we required the sample size to be a multiple of 6 (i.e., number of possible combinations). Therefore, our target sample size was 24 participants. In accordance with our preregistration, participants were excluded and replaced if they failed to learn or accurately perform the motor and object sequences (criteria detailed in the behavioral analysis section), did not adhere to experimental instructions (e.g., maintaining a regular sleep schedule or refraining from drug and alcohol use; see procedure), or did not obtain sufficient sleep to receive adequate stimulation (i.e., less than 50 stimuli).

A total of 32 young (aged 18-35) healthy volunteers (13 female, mean age=24, SD=4.8) were recruited for this study through local advertisements posted on the University of Utah campus. Inclusion criteria were verified with online screening questionnaires and included: 1) no previous extensive training with a musical instrument requiring dexterous finger movements (e.g., piano, guitar) or as a professional typist, 2) free of medical, neurological, psychological, or psychiatric conditions, including depression and anxiety as assessed by the Beck’s Depression and Anxiety Inventories (Beck et al. 1961, 1988), 3) no indications of abnormal sleep, as assessed by the Pittsburgh Sleep Quality Index (PSQI; Buysse et al. 1989), 4) not considered extreme morning or evening types, as quantified with the Horne & Ostberg chronotype questionnaire (Horne and Ostberg 1976); and, 4) free of psychoactive and sleep-influencing medications. We also assessed handedness using the Edinburgh Handedness Inventory (Oldfield, 1971) and daytime sleepiness with the Epworth sleepiness scale (Johns 1991). All study procedures were approved by the University of Utah institutional review board (IRB_00155080). All participants provided written informed consent at the start of the study and received monetary compensation for their time. Out of the 32 participants recruited for this study, two participants withdrew after the habituation session as they were unable to continue due to scheduling issues. Furthermore, six participants were excluded as they presented insufficient sleep duration to receive the pre-registered minimum number of stimulations (two participants excluded after experimental session 1 and four after experimental session 2). Twenty-four participants were eventually included in the final study sample. Participant demographics as well as information on their sleep and vigilance are presented in the Supplemental Tables S5-S6.

### Task

All participants performed an adapted version of the explicit Serial Reaction Time Task (SRTT) previously used in our research to examine sequence learning in the two different memory domains (procedural vs. declarative) with the same task (Temudo et al., 2026, 2025). The task was designed such that participants learned a motor sequence and object sequence in two separate sessions. The task consisted of a 4-choice reaction time task in which participants were instructed to react to visual cues shown on a screen by pressing a key on a keyboard using their non-dominant hand. A large visual cue shown in the center of the screen corresponded to one of eight different images of objects that belonged to 1 of 4 object categories (animals, vehicles, fruit, and tools). The 8 different objects were constantly presented on the bottom of the screen in 2 rows of 4 objects, each mapped to the four fingers used to perform the task (i.e., no thumbs). Object cues were presented one after the other in the center of the screen and participants were instructed to respond to the cue as fast and as accurately as possible by pressing the corresponding key according to the object/key mapping displayed at the bottom of the screen (Figure 1A). Importantly, the key-object mapping displayed on the bottom of the screen changed after each stream of 8 elements (i.e., every 8 objects / 8 key presses) in order to orthogonalize motor and object sequence learning within each learning session. With this manipulation, we developed three different versions of the task (Temudo et al., 2026, 2025). In the ***random condition***, the series of both responses and objects followed a random order. Performance improvement in this condition is expected to be minimal and represents general visuomotor habituation to the task, given that there is no (object or motor) sequence to learn. This task condition was administered at the start of each experimental session to measure baseline performance levels (see experimental procedure and Figure 1A). In the ***motor sequence condition***, the responses on the keyboard followed a repetitive sequential pattern (motor sequence, e.g., 3,4,1,2,1,3,4,2, whereby 1 and 4 represent the little and index finger, respectively) while the stream of objects on the screen followed a random order. In this task, performance improvement (as compared to random) therefore specifically reflects the learning of the sequence of movements. This task condition was learned during the first experimental session (see experimental procedure and Figure 1A). In the ***object sequence condition***, the stream of objects on the screen followed a sequential pattern (object sequence, e.g., car, saw, pineapple, giraffe, banana, plane, axe, elephant) while the responses on the keyboard followed a random order. Improvement in performance on this task (compared to random) is therefore specifically related to the knowledge of the series of objects. This task condition was learned during the second experimental session (see experimental procedure and Figure 1A). Altogether, this task allows us, by contrasting performance between the random and sequential SRTT within each experimental session, to extract sequence-specific learning in both memory domains (i.e., beyond general visuomotor learning). As in our previous TMR research (Nicolas et al. 2022), participants learned 2 sequences within each learning session (e.g., 2 motor sequences within the motor learning session). This design allows one sequence to be reactivated during sleep, while the other serves as a no-reactivation control condition. Practice was organized in blocks consisting of four repetitions of one of the sequences with a 1.5s interval between repetitions, then, after a 2s interval, four repetitions of the other sequence. For all task variants, each block included 64 key presses separated by 10s rest periods. To ensure clear differentiation between the two object sequences, a different set of 8 objects was used for the two different sequences (see below for example). Furthermore, each sequence was associated with a specific sound that was played at the start of each sequence repetition during task practice (see acoustic stimuli for details). The response stimulus interval (RSI) was 1 s. During this interval, the 8 objects on the bottom remined on the screen while a green fixation cross in the center of the screen replaced the central cue object. The longer 10 s rest periods as well as the intervals between sequence repetitions were indicated with a red fixation cross in the center of the screen while red frames replaced the locations of the 8 objects that displayed the key-object mapping at the bottom of the screen. Participants were made aware during the experimental visits that the order of objects/movements followed a repeating sequential pattern but were not told the precise sequence or the number of elements.

As declarative memory is defined as the conscious recollection of previous events, we implemented an **object generation/recall test** similar as in earlier research that asked participants to reconstruct the temporal order of the object items (Hsieh et al., 2014). For completeness, **a motor generation task** (King et al., 2019) was also implemented to test for any explicit knowledge of the learned sequences of movements. These tasks were administered at the end of each learning session to also ensure that participants associated each sound to the corresponding sequence. In the generation tasks, participants were presented with auditory cues corresponding to the two sequences learned during the session (3 repetitions for each sequence) and were instructed to reproduce the associated sequence either verbally (object sequence task) or via finger movements (motor sequence task). Participants therefore generated the two object sequences after the object sequence learning session and the two motor sequences after the motor sequence learning session (see procedure and Figure 2A). Responses were self-paced, with instructions to prioritize accuracy over speed.

Importantly, this paradigm also allowed us to create higher-order associations between memory domains (i.e., between the different learning session, see experimental procedure and Figure 1B). Specifically, the task was designed such that there were - or not - associations between fingers and object categories across the motor and object sequence learning tasks learned in the two separate sessions. Sequences were created based on the key-object mapping rule shown in the left panel of Figure 1B (key 1: fruit; key 2: animal, key 3: vehicle and key 4: tool). Note that within each learning session on day 1, object and motor items are orthogonalized as described above; therefore, associations between memory domains are only performed *across* learning sessions (see color coding across sessions in Figure 1B) and not within each learning session. In order to accommodate for the nature of the design (within-subject design with 2 learned sequences [reactivated and not-reactivated] per memory domain / session), 6 different motor sequences (seq A: 3,4,1,2,1,3,4,2; seq B: 4,1,2,3,3,2,4,1; seq C: 2,4,1,3,4,3,1,2; seq D: 4,3,2,3,4,1,1,2; seq E: 1,3,1,4,2,3,4,2; seq F: 2,4,4,2,1,3,1,3) and 6 corresponding object sequences (seq A: car, saw pineapple, giraffe, banana, plane, axe, elephant; seq B: hammer, pear, camel, bus, helicopter, zebra, screwdriver, orange; seq C: elephant, axe, banana, plane, saw, car, pineapple, giraffe; seq D: screwdriver, helicopter, zebra, bus, hammer, orange, pear, camel; seq E: pineapple, plane, banana, saw, elephant, car, axe, giraffe; seq F: camel, screwdriver, hammer, zebra, pear, helicopter, orange, bus) were designed. In the ***associated visit***, each object sequence was paired with its corresponding motor sequence (e.g., Motor sequence A paired with the associated Object sequence A, see supplementary Table S1 for all pairs used) such that each finger in the motor sequence was associated to a particular object category in the object sequence (e.g. finger 4 in the motor sequence was always associated to tools in the object sequence; see color coding in Figure 1B *Associated*). In contrast, in the ***unassociated visit***, each object sequence was paired with a different motor sequence (e.g., Motor sequence A paired with the unassociated Object sequence C, supplementary table S1 for all pairs used) such that each finger in the motor sequence was associated to objects from different categories in the object sequence (e.g. finger 4 in the motor sequence was associated to a vehicle and a fruit; see color coding in Figure 1B *Unassociated*). Consequently, motor and object sequences did not share such a higher-order association between the two learning sessions. In this within-subject design, participants therefore learned during the ***associated visit***, 2 (out of 6) pairs of associated sequences, while during the ***unassociated visit***, the 4 other pairs were shuffled in order to create 2 pairs of sequences that were unassociated (see supplementary Table S1 for all combination of pairs used). Pair combinations were counterbalanced across visits (associated and unassociated) and participants. Importantly participants were not explicitly informed that the motor and object tasks shared an underlying sequential structure and were not instructed to look for or identify the correspondence between the two sequences. A debriefing questionnaire administered at the end of the experiment indicated that none of the participants (0/24) reported any awareness of potential relationships between the tasks which suggest that participants did not develop explicit awareness of the shared structure between the different tasks during the associated visit.

### Acoustic stimuli

During each experimental visit (associated and unassociated visits, see procedure below), a specific sound was associated to each of the 4 learned sequences (2 motor and 2 object sequences). During each experimental nap, one of the 4 sounds linked to one object memory trace (memory cue) was replayed in an alternating fashion with a control sound that was not related to learning (control cue). A total of 10 sounds was therefore required for this design (2 linked with each object sequence, 2 linked with each motor sequence and 1 control sound in each of the associated and unassociated visits, see procedure below). The 10 sounds (bleep, cowbell, crack, swoosh, keys, laser, pop, snare, sparkle and zip) were taken from https://freesound.org, and were edited to each last 500ms. Sounds were pseudo-randomly assigned to the different conditions (motor and object sequences, reactivated vs. not reactivated conditions, memory vs. control cue) across visits and individuals. At the start of the experimental visit, auditory detection thresholds were determined by the participants themselves using a transformed 1-down 1-up procedure (Levitt 1971; Leek 2001) separately for each of the 5 sounds used in the specific visit. Subsequently, the sound pressure level was set to 8000% of the individual auditory threshold during the learning sessions and to 300% for auditory stimulation during sleep, to limit the risk of awakening during the nap. During both task practice and sleep, sounds were played to participants via in-ear earphones.

### Procedure

This study employed a within-subject design (Figure 2A). Approximately one week prior to the experimental sessions, participants were first invited to complete a habituation session. During this session, they performed 20 blocks of the random condition of the SRTT to exhaust general – i.e., non-sequence-specific – enhancement in performance on the task and minimize order (transfer) effects during the subsequent experimental sessions with the sequential task. They were also invited to take a 90-minute habituation nap that was monitored with polysomnography (see below). Participants then completed two experimental visits one week apart, each involving either the **associated** version or the **unassociated** version of the paradigm (order counterbalanced across individuals). Participants were asked to respect a constant sleep schedule during the 3 days/nights preceding each experimental visit (according to their own schedule ±1h; latest bedtime: 1am; minimum 7h of sleep; no naps) and compliance with this schedule was assessed with both sleep diary and wrist actigraphy (ActiGraph wGT3X-BT, Pensacola, FL; see Table S6).

Each experimental visit began in the morning (between 8am-10am) and lasted approximately 6 hours. First, participants completed a brief questionnaire to collect information regarding their sleep patterns in the previous 24 hours (St Mary’s Hospital Sleep Questionnaire (SMS); Ellis et al., 1981), their subjective feelings of alertness (Stanford Sleepiness Scale (SSS); (MacLean et al., 1992), and any consumption of alcohol and/or recreational drugs in the last 24 hours. Next, participants performed a psychomotor vigilance task (PVT) to obtain an objective measure of their alertness. For the PVT, participants were asked to fixate on a fixation cross in the center of the screen and to press the space bar as fast as possible when the fixation cross disappeared. A timer appeared on the screen displaying the participants’ reaction time in milliseconds. Participants completed 50 trials of the PVT. Next, the participants practiced 4 blocks of the random condition to measure baseline performance. This was then followed by the auditory threshold task (see acoustic stimulation section) before the training on the sequential SRTT. For the sequential SRTT, participants first learned 2 motor sequences with 20 blocks of practice. Each motor sequence was associated with a different auditory stimulus which was played through earphones and cued at the start of each sequence repetition. After a short break offered to minimize the confounding effect of fatigue (Pan and Rickard 2015), performance on the 2 motor sequences was tested with 4 additional blocks of practice which was then followed by the auditory-cued motor generation task. After a 15 min break, participants next learned 2 object sequences with 20 blocks of practice. Each object sequence was associated with a different auditory stimulus played at the start of each sequence repetition. Performance on the 2 object sequences was also tested with 4 additional blocks of practice that was followed by an auditory-cued object generation task. Note that the order of the motor and object sequence learning sessions was not counterbalanced across participants and that object sequence learning always followed motor sequence learning. This design choice was based on our previous work (Temudo et al., 2026) suggesting that interactions between memory domains are stronger when object learning followed motor learning. To further enhance these interactions, the interval between the two learning sessions was kept short (15 min, Mosha & Robertson, 2016).

After the training, participants were invited to take a break and were provided with a standardized lunch. They were then equipped with the EEG system (see details in Polysomnography and TMR protocol section) and the earphones and invited to take a 90-minute nap in the sleep laboratory. The 4 sounds corresponding to all learned sequences during the visit and an additional 5^th^ control sound (not associated to learning) were played at 300% of the perception threshold before the nap to verify that the participants could perceive the sounds. Participants were instructed that they may or may not receive auditory stimulation during sleep. When the participants reached NREM sleep during the nap, one of the sounds linked to one of the object sequences (*reactivated object memory*) was replayed. The sound linked to the other object sequence was not replayed and this sequence therefore served as a *not reactivated control condition*. None of the sounds linked to the motor task were played during the nap. With this design, our goal was to indirectly reactivate a motor memory trace via higher-order associations with the reactivated object memory trace (associated visit; *motor memory reactivated by association*, see Figure 2B). During the unassociated experimental visit, we did not expect the motor memory to be indirectly reactivated as there was no higher-order associations with the reactivated object (unassociated visit; *motor memory not reactivated*). A control sound that was not linked to any memory trace (control cue) was also played during the nap and served as a control for our EEG analyses (to measure brain responses elicited by a sound that did not carry memory content (Nicolas et al., 2022)). We hypothesized that the reactivation process during the associated visit would involve a higher memory load (two reactivated memories: *object reactivated* and *motor reactivated by association*) than the unassociated visit during only one memory was reactivated (*object reactivated*). At the end of the nap, the EEG system was removed, and a retest session was administered after a minimum of 30 min to allow for sleep inertia to dissipate. During the retest session, participants completed 4 blocks of each of the motor sequences, followed by 4 blocks of each of the object sequences followed by 4 blocks of the random sequence to assess sequence-specific learning. Auditory tones were also played during task retest as during the training sessions. Approximately one week later, participants were invited back for a second experimental visit which followed the same procedure as the first experimental visit but with the other association condition (associated vs. unassociated). Visit order (associated or unassociated) was counterbalanced across individuals. The combination between sequences (A-F), sounds (1-10), task conditions (motor vs. object), reactivation conditions (reactivated vs. not-reactivated) and cue conditions (memory vs. control cues) was pseudorandomized between participants.

### Polysomnography and TMR protocol

The habituation and experimental naps were monitored with a wireless digital sleep recorder (LiveAmp, Brain Products, Gilching, Germany; bandwidth: DC to Nyquist frequency) and digitized at a sampling rate of 1000 Hz. Standard electroencephalographic (EEG) recordings were made from 11 electrodes (FPz, Fz, Cz, Pz, Oz, F3, F4, C3, C4, P3, P4) according to the international 10–20 system. A2 was used as the recording reference and A1 as a supplemental individual EEG channel. The recording ground consisted of an electrode placed on the middle of the forehead. Bipolar vertical and horizontal eye movements (electro-oculogram, EOG) were recorded from electrodes placed above and below the right eye and on the outer canthus of both eyes, respectively. Bipolar submental electromyogram (EMG) recordings were made from the chin. Electrical noise was filtered using a 60-Hz notch. Impedance was kept to < 5kΩ for all electrodes. During the habituation nap, only FPz, Fz, Cz, Pz, Oz, horizontal and vertical EOGs and EMGs were set-up. Note that due to experimental error, one experimental session (unassociated visit) of one participant was recorded with this subset of electrodes (i.e., with the habituation montage) instead of the 11 electrodes mentioned above.

During the experimental nap, TMR was applied during NREM 2–3 sleep stages. To do so, sleep stages were visually scored online by a researcher (AT) monitoring the PSG recordings (following guidelines from the American Academy of Sleep Medicine (Iber, 2007)). For online sleep scoring, PSG recordings were displayed using 30 s long epochs with EEG and EOG data filtered from 0.5 to 30 Hz and EMG data filtered between 20 and 200 Hz. Auditory cues corresponding to one object sequence (directly reactivated object memory) as well as the control cue not carrying memory content were sent during NREM 2–3 sleep stages. Each type of auditory cue (memory and control cues) was presented in alternating 3-min-long stimulation blocks that were separated by 1 min of silence / rest blocks (see Figure 2C). During the stimulation blocks, auditory tones were played every 5 s. Auditory stimulation was paused if REM, NREM stage 1 or wakefulness were detected.

### Behavioral analyses

#### Performance measures

During performance of the SRT tasks, the keys pressed and timing of the key presses were recorded, and performance was quantified using speed and accuracy measures. Specifically, performance speed was defined as the time between the visual cue onset and the corresponding response for the correct trials. For accuracy, a trial was classified as “correct” if the key pressed by the participants matched the object cue and as “incorrect” if it did not. The averaged response times for correct key presses and the percentage of correct key presses were computed for each block of practice for each version of the task. Individual trials (i.e., responses) were excluded from the analyses if the measured response time was greater than 3 standard deviations above or below the participant’s mean response time for that block [1.53% trials excluded in the associated visit and 1.58% excluded in the unassociated visit, t(23) = -.48, p = .63].

Performance from the 4 test blocks at the end of training were averaged to calculate pre-TMR performance for each task condition and learned sequence, and performance from the 4 retest blocks after the nap were averaged to calculate post-TMR performance for each sequence and each task condition. Offline changes in performance reflecting the consolidation process were computed as the difference between pre- and post-TMR performance for each of the 4 sequences. A TMR index for the object condition was calculated as the difference in offline changes in performance between the reactivated and not reactivated conditions for each visit separately (associated/unassociated). The TMR index for the motor condition was calculated as the difference in offline changes in performance between the indirectly reactivated and the not indirectly reactivated conditions for each visit separately (associated/unassociated). For the generation tasks, the percentage of items (keys or object) placed in their correct ordinal position as well as the percentage of correct transitions (i.e., two correct consecutive items) were computed for each sequence, condition and visit, as the average across the three attempts.

#### Confirmatory analysis

Our primary confirmatory analysis consisted in examining whether memories linked through higher-order associations would benefit from TMR more than unassociated memories. We therefore analyzed offline changes in performance (speed and accuracy) using a pre-registered repeated-measures ANOVA using task (motor vs. object), reactivation condition (reactivated vs. not reactivated) and experimental visit (associated vs. unassociated) as within-subject factors.

#### Negative control analyses

To confirm that any observed differences in the results of our confirmatory analysis could be attributed to our experimental manipulation, a series of control analyses were conducted on the training data. First, baseline motor execution was compared using separate repeated-measures ANOVAs on performance (speed and accuracy) of the random serial reaction time task with visit (associated vs. non-associated), task (object vs. motor) and block (1-4) as within subject factors. Results for baseline execution indicated comparable baseline performance across visits and task conditions and can be found in the Section 2 of the Supplementary text. Second, performance (speed and accuracy) on the sequential SRTT during initial learning (separately for training and test blocks) was analyzed using repeated-measures ANOVAs using task (motor vs. object), visit (associated vs. non-associated), reactivation condition (reactivated vs. non-reactivated) and block (20 for training, 4 for test) as within-subject factors. These analyses assessed whether there was 1) equivalent baseline performance between the reactivated- and non-reactivated conditions for each task and visit, 2) equivalent performance between experimental visits for each task and 3) equivalent changes in performance (interference/transfer) between tasks and between experimental visits. Corresponding results are presented in the Supplemental Section 3 and collectively indicate that performance speed and accuracy did not differ between reactivated and non-reactivated sequences across tasks and visits. Finally, sequence awareness was compared using repeated-measures ANOVAs on measures extracted from the generation task (i.e., percentage of items in their correct ordinal position and percentage correct transitions) with visit (associated vs. non-associated), task (object vs. motor) and reactivation condition (reactivated vs. non-reactivated) as within-subject factors. Results of the generation task are presented in the Supplemental Section 7 and indicated that, as expected, participants developed explicit knowledge of the series of objects and movements.

### EEG analyses

#### Offine sleep scoring

Offline sleep stage scoring was completed by a certified sleep technologist blind to the research question, protocol and stimulation periods/conditions and according to the criteria of the American Academy of Sleep Medicine (Iber et al., 2007). Scoring was conducted using the *Counting Sheep PSG* toolbox (https://github.com/stuartfogel/CountingSheepPSG) (Ray et al., 2024). Data was visually scored in 30 s epochs and band pass filters were applied between 0.3 and 35 Hz for EEG and EOG signals, and 10 and 50 Hz for EMG. The duration and latency of each sleep stage, as well as stimulation accuracy (i.e., % of auditory cues sent during NREM2-3 sleep), total sleep time and sleep efficiency (% time asleep in NREM2-3 and REM sleep relative to total time allowed to sleep) were extracted from the scored dataset and compared between experimental visits using paired t-tests (see Table S2-S3).

#### EEG preprocessing

EEG data preprocessing was carried out using functions supplied by the fieldtrip toolbox (Oostenveld et al. 2011). Specifically, data were re-referenced to the average of channels A1 and A2 and band-pass filtered between 0.1 and 30 Hz. Thirty-second epochs were visually inspected, and segments contaminated by muscle activity or eye movements were rejected. Additional artifacts were removed using independent component analysis.

#### Sleep event detection

Preprocessed cleaned data was downsampled to 500 Hz and transferred to the python environment. SOs and spindles were detected automatically in NREM2-3 sleep epochs on all the channels, by using algorithms implemented in the YASA open-source Python toolbox (Vallat and Walker 2021). Regarding SO detection, the algorithm used was a custom adaptation of (Massimini et al. 2004; Carrier et al. 2011). Specifically, the data were filtered between 0.3 and 2 Hz using a finite impulse response (FIR) filter with a 0.2 Hz transition band, resulting in –6 dB cutoff points at 0.2 and 2.1 Hz. Negative peaks with amplitudes between –40 and –200 μV and positive peaks between 10 and 150 μV were first detected in the filtered signal. Negative peaks were then paired with subsequent positive peaks, after which a set of criteria was applied to identify SOs: (1) negative peak duration between 0.3 and 1.5 s, (2) positive peak duration between 0.1 and 1 s, (3) negative peak amplitude between 40 and 300 μV, (4) positive peak amplitude between 10 and 200 μV, and (5) peak-to-peak (PTP) amplitude between 75 and 500 μV.

Spindle detection was based on an algorithm inspired by the A7 method described by Lacourse et al. (2019). Specifically, relative power in the spindle frequency band (8–18 Hz) was first computed with respect to total broadband power (1–30 Hz) using Short-Time Fourier Transforms with 2-s windows and 200 ms overlap. Next, the algorithm calculated the moving root mean square (RMS) of the EEG signal filtered in the sigma band using a 300 ms window with a 100 ms step size. A moving correlation between the broadband signal (1–30 Hz) and the EEG signal filtered in the spindle band was then computed. Sleep spindles were detected when three criteria were simultaneously met: (1) relative sigma power exceeded 0.2, (2) the moving RMS surpassed a threshold of RMS_mean + 1.5 × RMS_SD, and (3) the moving correlation exceeded 0.65. Detected events shorter than 0.3 s or longer than 3 s were discarded. Spindles detected across different channels within 500 ms were considered to reflect the same event and were therefore merged.

SOs and spindles were detected during the stimulation intervals of both memory-cue and control-cue conditions for both visits (associated and unassociated) as well as during rest intervals (i.e., NREM 2–3 epochs without auditory stimulation). Note that, due to experimental error, stimulation triggers were not recorded for one participant during one experimental session (unassociated visit) and the sleep metrics mentioned above could therefore not be extracted for that session (associated visit: N=24; unassociated visit N=23). With respect to the detected SOs, we extracted for each participant, each stimulation condition and channel, *SO density*, i.e., the number of SOs per total time in minutes spent in stimulation or rest intervals (and see ERP for *SO amplitude* analyses). Concerning spindles, we extracted for each participant, stimulation condition and channel, *spindle density* (i.e., the number of spindles per total time in minutes spent in stimulation or rest intervals), *spindle amplitude* (computed as the peak-to-peak amplitude (µV) in the sigma-filtered data) and *spindle frequency*. These different SO and spindle features were then averaged across channels and compared between stimulation conditions (memory-cue, control-cue, no stimulation) and visits (associated vs. unassociated) using repeated-measures ANOVAs (results presented in the Supplemental Section 4 and Figure S5).

#### Event-related analyses

##### Events of interest

Event-related analyses were performed using either auditory cues or the detected SOs as events (associated visit: N=24; unassociated visit N=23). *Auditory-cue events* were obtained by segmenting the data into epochs time-locked to auditory cue onset (from –1 to 3 s relative to auditory cue onset #, see section 6 of Supplemental information) for each condition (associated memory cue, associated control cue, unassociated memory cue, unassociated control cue). The average number of artifact-free auditory cues used in these analyses was 176.33 [95% CI: 157.54-195.13] and 172.29 [95% CI: 151.51-193.07] for the memory and control cues respectively in the associated visit and 177.22 [95% CI: 154.79-199.64] and 175.39 [95% CI: 154.15-196.63] for the memory and control cues respectively in the unassociated visit. The number of auditory cues did not differ between cue and visit conditions (visit: F(1,22) = .31, p = .58, ηp² = .01; cue condition: F(1,22) = 1.10, p = .31, ηp² = .05; visit x cue condition: F(1,22) = .23, p = .63, ηp² = .01).

*SO events* were obtained by segmenting the data into epochs time-locked to the trough of the detected SOs (from –2 to 2 s relative to the trough of the SO). SO events were extracted from 3 different intervals, i.e., periods of memory-cue stimulation, control-cue stimulation and no-stimulation (i.e., any NREM2-3 sleep period during which there was no auditory stimulation). Table S4 in the supplementary information presents the number of SO detected on each EEG channel. The average number of artifact-free SO trials included in these analyses was 114.83 [95% CI: 74.20-155.45], 117.21 [95% CI: 74.56-159.85] and 140.42 [95% CI: 64.71-216.13] for the memory-cue, control-cue and rest periods respectively in the associated visit and 98.59 [95% CI: 66.97-130.22], 106.18 [95% CI: 74.06-138.30] and 102.52 [95% CI: 64.64-140.40] for the memory-cue, control-cue and rest periods respectively in the unassociated visit. The number of detected SOs included in the analyses did not differ between intervals and visits (visit: F(1,22) = .97, p = .34, ηp² = .04; interval: F(2,44) = 1.19, p = .32, ηp² = .05; visit x interval: F(2,44) = .80, p = .46, ηp² = .04).

##### Potential analyses

Event-related potential (ERP) analyses using either auditory cues or SOs as events were performed using the fieldtrip toolbox (Oostenveld et al. 2011) on down sampled data (100 Hz).

*Auditory-evoked responses* were obtained by averaging all trials in each cue condition separately (memory and control cues) during the associated and unassociated visits. Individual ERPs computed on each channel were then baseline corrected by subtracting the mean amplitude from –0.3 to –0.1 s relative to cue onset # (see section 6 in the Supplemental information). To assess cue-related effects within each visit, we first computed ERP difference maps contrasting memory and control cues separately for the associated and unassociated visits. These contrasts were then subtracted to obtain the ERP difference between visits ([memory – control] x [associated – unassociated]). Spatio-temporal clusters in which ERP amplitude was significantly different between conditions (and visits) were identified using cluster-based permutation (CBP) tests (Maris and Oostenveld 2007, see below) on a 0 to 2.5 s window relative to cue onset.

*SO-locked responses* were extracted with a similar procedure as above but time-locked to the trough of detected SOs. SO-locked potentials were obtained by averaging all SOs in each interval, i.e., for each cue condition and during rest periods separately (memory-cue stimulation, control-cue stimulation and no-stimulation during the associated and unassociated visits). Significant spatio-temporal clusters were identified with CBP tests performed on a –1.5 to 1.5 s window relative to the SO trough (3 pairwise comparisons: memory – control, memory – rest, control – rest in the associated and unassociated visits separately and between visits).

##### Time-frequency analyses

To examine oscillatory activity induced by the auditory stimulation and the detected SOs, time–frequency representations (TFRs) of power were computed for each experimental condition and channel. For both the auditory- and the SO-locked analyses, power was estimated between 5 and 30 Hz using a Hanning taper and fast Fourier transform (FFT) with an adaptive sliding time window of five cycles per frequency (Δt = 5/f; step size = 20 ms). Individual TFRs were then converted to baseline-relative power change using a baseline period from –0.3 to –0.1 s relative to cue onset # for the auditory-cue-locked analyses and –2 to 2 s relative to SO trough for the SO-locked analyses. As in the ERP analysis, TFRs were first compared within each visit by contrasting the cue condition (i.e., memory and control cues) for the auditory-locked analyses and the intervals (i.e., memory-cue vs. control-cue vs. rest periods) for the SO-locked analyses, and then between visits (associated vs. unassociated). Spatio-temporal-frequency clusters were identified using cluster-based permutation (CBP) tests conducted in a 0 to 2.5 s time window relative to auditory-cue onset for the auditory-locked analyses and a –1.5 to 1.5 s time window relative to SO trough for the SO-locked analyses. For both analyses, the frequency window was 5–30 Hz and the spatial dimension included all channels.

##### Brain-behavior correlation analyses

We performed correlation analyses between the TMR index (for both object and motor tasks separately for each experimental visit; see performance measures above) and the following EEG-derived data extracted from each experimental visit: the memory-vs. control-cue difference in (1) auditory-locked ERP amplitude, (2) SO-locked amplitude, (3) auditory-locked power and (4) SO-locked power. Significant spatio-temporal and spatio-temporal-frequency clusters were identified using CBP tests for the potential and power outcomes, respectively.

##### Statistical analyses

For both auditory-cue- and SO-trough-locked analyses, ERP and TF data were analyzed using nonparametric cluster-based permutation (CBP) tests (Maris and Oostenveld 2007) implemented in the FieldTrip toolbox. Paired *t*-tests were used to compare stimulation conditions and/or visits, with cluster-based correction applied to control for multiple comparisons across time and space for ERP analyses and for time, frequency, and space for TF analyses. Samples exceeding a cluster-forming threshold of α = 0.01 were grouped into clusters based on adjacency (time–space for ERP; time–frequency–space for TF). For each cluster, the sum of *t*-values—referred to as the *cluster size*—was used as the test statistic. Cluster sizes were then compared against a null distribution generated from 500 permutations with randomly shuffled condition labels within participants. Clusters were considered significant at *p* < 0.05. Cohen’s *d* is reported for CBP contrasts.

## Author contributions

GA, AT C BRK designed the experiments. AT conducted the experiments. GA, AT, BRK C JN contributed to acquisition and analytic tools. AT C GA analyzed the data. All authors contributed to the manuscript.

## Acknowledgement

We thank Elise Perry, Krista Hamilton and Ashmita Karki for their assistance with data collection.

