## Supplementary for "Cross-domain competition for memory reactivation resources during sleep"

**This PDF file includes:**

Supporting text 1-7
Figures S1 to S5
Tables S1 to S5

1. Offline changes in performance

#### A. Speed

A repeated-measures ANOVA was conducted on changes in performance speed (post-nap – pre-nap) with visit (associated vs. unassociated), task (motor vs. object), and reactivation condition (sequence reactivated vs. non-reactivated) as within-subject factors (Figure 3B in the main text). While the main results of this analysis are presented in the main text, the results of the full statistical model are reported here. Results revealed that the motor task was performed faster than the object task ($F(1,23) = 9.24$, $p = .006$, $\eta p^2 = .29$). A significant task × reactivation condition interaction ($F(1,23) = 4.22$, $p = .05$, $\eta p^2 = .16$) was also observed but contrary to our expectations, it reflected greater gains for the non-reactivated sequence compared to the reactivated sequence in the object task (reactivated vs. non reactivated: $F(1,23) = 3.48$, $p = .08$, $\eta p^2 = .13$) while no such effect was observed for the motor task (reactivated vs. non reactivated: $F(1,23) = 0.71$, $p = .41$, $\eta p^2 = .03$). Also in contrast to our hypotheses, there was no main effect of visit ($F(1,23) = 0.31$, $p = .58$, $\eta p^2 = .01$) or reactivation condition ($F(1,23) = 0.53$, $p = .47$, $\eta p^2 = .02$), nor were there significant interactions involving visit (visit × task: $F(1,23) = 1.67$, $p = .21$, $\eta p^2 = .07$; visit × reactivation: $F(1,23) = 1.50$, $p = .23$, $\eta p^2 = .06$; visit × task × reactivation: $F(1,23) = 0.06$, $p = .81$, $\eta p^2 = .003$).

#### B. Accuracy

A repeated-measures ANOVA was conducted on changes in performance accuracy (post-nap – pre-nap) with visit (associated vs. unassociated), task (motor vs. object), and reactivation condition (sequence reactivated vs. non-reactivated) as within-subject factors (Figure S1). Results revealed that unlike performance speed, there was no significant main effect of task ($F(1,23) = .99$, $p = .33$, $\eta p^2 = .04$) nor a significant task × reactivation condition interaction ($F(1,23) = .19$, $p = .67$, $\eta p^2 = .008$). There was also no main effect of visit ($F(1,23) = .12$, $p = .73$, $\eta p^2 = .005$) or reactivation ($F(1,23) = 3.15$, $p = .09$, $\eta p^2 = .12$), or any significant interactions with visit (visit × task: $F(1,23) = 1.36$, $p = .26$, $\eta p^2 = .06$; visit × reactivation: $F(1,23) = 1.28$, $p = .27$, $\eta p^2 = .05$; visit × task × reactivation: $F(1,23) = .40$, $p = .53$, $\eta p^2 = .02$). These results indicate that both TMR and higher-order associations between tasks did not influence changes in performance accuracy.

### 2. Assessment of baseline performance on random SRTT

#### A. Speed

Baseline motor execution was assessed using the four blocks of the random task performed at the start of each training session (motor and object) and visit (associated vs. unassociated). A repeated-measures ANOVA with block (1–4), visit and task as within subject factors revealed that performance speed remained stable across blocks ($F(3,69) = 1.56$, $p = .21$, $\eta p^2 = .06$). Importantly, there was no main effect of visit (visit: $F(1,23) = 1.31$, $p = .26$, $\eta p^2 = .05$; visit x block: $F(3,69) = .33$, $p = .80$, $\eta p^2 = .01$) or a main effect of task (task: $F(1.23) = .09$, $p = .76$, $\eta p^2 = .004$; task x block: $F(3,69) = 1.49$, $p = .23$, $\eta p^2 = .06$) or any significant interactions between these factors (visit x task: $F(1,23) = .00$, $p = .97$, $\eta p^2 = .00$; visit x task x block $F(3,69) = .73$, $p = .54$, $\eta p^2 = .03$). These results indicate comparable baseline performance speed across visits and task conditions.

#### B. Accuracy

The repeated-measures ANOVA on performance accuracy with block (1–4), visit and task as within subject factors revealed that performance accuracy fluctuated across blocks ($F(3,69) = 3.12$, $p = .03$, $\eta p^2 = .12$). Importantly, there was no main effect of visit (visit: $F(1,23) = .91$, $p = .35$, $\eta p^2 = .04$; visit x block: $F(3,69) = 1.63$, $p = .19$, $\eta p^2 = .07$) or a main effect of task (task: $F(1.23) = 2.48$, $p = .13$, $\eta p^2 = .10$; task x block: $F(3,69) = .95$, $p = .42$, $\eta p^2 = .04$) or any significant interactions between these factors (visit x task: $F(1,23) = 0.32$, $p = .58$, $\eta p^2 = .01$; visit x task x blocks $F(3,69) = .20$, $p = .89$, $\eta p^2 = .009$). These results indicate comparable baseline performance accuracy across visits and task conditions.

### 3. Performance during initial training

#### A. Speed

We first verified that baseline performance (training and test) did not differ between reactivation conditions (reactivated vs. not-reactivated sequences) in the different task conditions (motor vs. object) and visits (associated vs. unassociated). The block x reactivation condition x task x visit repeated-measures ANOVA revealed a significant block effect during training (block: $F(19,437) = 40.27$, $p < .001$, $\eta p^2 = .64$) that did not interact with any other factors (visit x block: $F(19,437) = .81$, $p = .70$, $\eta p^2 = .03$; task x block: $F(19,437) = 1.50$, $p = .08$, $\eta p^2 = .06$; reactivation x block: $F(19,437) = .83$, $p = .68$, $\eta p^2 = .04$; visit x task x block: $F(19,437) = 1.02$, $p = .44$, $\eta p^2 = .04$; visit x reactivation x block: $F(19,437) = 1.72$, $p = .03$, $\eta p^2 = .07$; task x reactivation x block: $F(19, 437) = 1.23$, $p = .23$, $\eta p^2 = .05$; visit x reactivation x task x block: $F(19,437) = 1.45$, $p = .10$, $\eta p^2 = .06$) suggesting that block-to-block performance improvements during learning was similar between reactivation conditions, task and visits (Figure 3A in main text). Performance stabilized during test (block: $F(3,69) = .60$, $p = .62$, $\eta p^2 = .03$) similarly between reactivation and task as well as between visits (visit x block: $F(3,69) = .36$, $p = .78$, $\eta p^2 = .02$; task x block: $F(3,69) = 1.84$, $p = .15$, $\eta p^2 = .07$; reactivation x block: $F(3,69) = 1.95$, $p = .13$, $\eta p^2 = .08$; visit x task x block: $F(3,69) = .18$, $p = .91$, $\eta p^2 = .008$; visit x reactivation x block: $F(3,69) = 1.24$, $p = .30$, $\eta p^2 = .05$; task x reactivation x block: $F(3,69) = 1.13$, $p = .34$, $\eta p^2 = .05$; visit x reactivation x task x block: $F(3,69) = 2.45$, $p = .07$, $\eta p^2 = .10$). Results also revealed a significant effect of task (training: $F(1,23) = 22.14$, $p < .001$, $\eta p^2 = .49$; test: $F(1,23) = 45.52$, $p < .001$, $\eta p^2 = .66$) whereby performance on the motor task was overall faster than on the object task but this effect did not interact with the reactivation condition or with the visit (training: task x reactivation: $F(1,23) = .08$, $p = .78$, $\eta p^2 = .004$; task x visit: $F(1,23) = .006$, $p = .94$, $\eta p^2 = .00$; task x visit x reactivation: $F(1,23) = .43$, $p = .52$, $\eta p^2 = .02$; test: task x reactivation: $F(1,23) = 3.58$, $p = .07$, $\eta p^2 = .14$; task x visit: $F(1,23) = .13$, $p = .72$, $\eta p^2 = .006$; task x visit x reactivation : $F(1,23) = .44$, $p = .51$, $\eta p^2 = .02$). Finally there was no significant effect of reactivation condition (training: $F(1,23) = .09$, $p = .76$, $\eta p^2 = .004$; test: $F(1,23) = .78$, $p = .39$, $\eta p^2 = .03$) or visit (training: $F(1,23) = 2.53$, $p = .13$, $\eta p^2 = .10$; test: $F(1,23) = .21$, $p = .66$, $\eta p^2 = .009$) or an interaction between these factors (training: $F(1,23) = .02$, $p = .89$, $\eta p^2 = .001$; test: $F(1,23) = .03$, $p = .86$, $\eta p^2 = .001$). Altogether, these results indicate that performance speed did not differ between reactivated and non-reactivated sequences across tasks and visits prior to sleep.

### B. Accuracy

The block x reactivation condition x task x visit repeated measures ANOVA on performance accuracy (Figure S1) revealed that accuracy remained stable across training blocks (block: $F(19,437) = 1.10$, $p = .34$, $\eta p^2 = .05$) and did not interact with any other factors (visit x block:

$F(19,437) = .89$, $p = .60$, $\eta p^2 = .04$; task x block: $F(19,437) = .91$, $p = .57$, $\eta p^2 = .04$; reactivation x block: $F(19,437) = 1.40$, $p = .12$, $\eta p^2 = .06$; visit x task x block: $F(19,437) = 1.21$, $p = .25$, $\eta p^2 = .05$; visit x reactivation x block: $F(19,437) = .76$, $p = .76$, $\eta p^2 = .03$; task x reactivation x block: $F(19, 437) = 1.28$, $p = .19$, $\eta p^2 = .05$; visit x reactivation x task x block: $F(19,437) = .43$, $p = .98$, $\eta p^2 = .02$) suggesting that block-to-block performance improvements during learning was similar between reactivation conditions, task and visits. Performance remained stable during test (block: $F(3,69) = 2.66$, $p = .06$, $\eta p^2 = .10$) similarly between reactivation condition and task as well as between visits (visit x block: $F(3,69) = 1.46$, $p = .23$, $\eta p^2 = .06$; task x block $F(3,69) = .23$, $p = .88$, $\eta p^2 = .01$; reactivation x block: $F(3,69) = .73$, $p = .54$, $\eta p^2 = .03$; visit x task x block: $F(3,69) = 1.68$, $p = .18$, $\eta p^2 = .07$; visit x reactivation x block: $F(3,69) = .24$, $p = .87$, $\eta p^2 = .01$; task x reactivation x block: $F(3,69) = .46$, $p = .71$, $\eta p^2 = .02$; visit x reactivation x task x block: $F(3,69) = .29$, $p = .83$, $\eta p^2 = .01$). Consistent with the speed results, accuracy also showed a significant effect of task (train: $F(1,23) = 26.12$, $p < .001$, $\eta p^2 = .53$; test: $F(1,23) = 13.17$, $p = .001$, $\eta p^2 = .36$) whereby participants were more accurate on the motor sequence task compared to the object sequence task, but this effect did not interact with the reactivation condition or with the visit (train: task x reactivation: $F(1,23) = .86$, $p = .36$, $\eta p^2 = .04$; task x visit: $F(1,23) = .49$, $p = .49$, $\eta p^2 = .02$¿ task x visit x reactivation: $F(1,23) = 1.04$ $p = .32$, $\eta p^2 = .04$; test: task x reactivation: $F(1,23) = .26$, $p = .61$, $\eta p^2 = .01$; condition x visit: $F(1,23) = .59$, $p = .45$, $\eta p^2 = .03$; task x visit x reactivation $F(1,23) = .24$, $p = .63$, $\eta p^2 = .01$). While no effect of reactivation condition was observed during training, this effect was significant during the last four test blocks (training: $F(1,23) = .004$, $p = .95$, $\eta p^2 = .00$; test: $F(1,23) = 6.17$, $p = .02$, $\eta p^2 = .21$), where accuracy was higher for the non-reactivated sequences compared to the sequences to be directly/indirectly reactivated. Finally, and importantly, there was no significant effect of visit ($F(1,23) = .07$, $p = .80$, $\eta p^2 = .003$; test: $F(1,23) = .12$, $p = .73$, $\eta p^2 = .005$) or an interaction between reactivation condition and visit (train: $F(1,23) = .02$, $p = .88$, $\eta p^2 = .001$; test: $F(1,23) = .60$, $p = .45$, $\eta p^2 = .03$). Altogether, these results indicate that performance accuracy did not differ between visits prior to sleep.

### 4. Sleep event characteristics

Sleep EEG data were analyzed to test whether memory (vs. control) cueing modulated the characteristics of electrophysiological markers critically involved in motor memory reactivation during sleep, i.e., SO and spindle events (Nicolas et al., 2022, 2025; Cairney et al., 2018). To do so, sleep spindles and SOs events were automatically identified during NREM2-3 sleep episodes in three different intervals referred to as memory cue, control cue and rest (no stimulation) using an established detection algorithm (Vallat & Walker, 2021) used in our previous research (e.g., Nicolas et al., 2022, 2025; see Supplementary Table S3 for spindle and SO counts for each channel, interval and visit). Auditory stimulation resulted in an increase in *SO density* (interval: $F(2, 44) = 46.27$, $p < .001$, $\eta p^2 = .68$) irrespective of the cue condition (Figure S6D; memory vs. control: $p = .12$; memory vs. rest: $p < .001$; control vs. rest: $p < .001$) or visit (visit: $F(1, 22) = 1.23$, $p = .28$, $\eta p^2 = .05$; visit x interval: $F(2, 44) = .63$, $p = .54$, $\eta p^2 = .03$). In contrast, spindle characteristics were modulated by the interval and visit conditions. Specifically, control cue stimulation resulted in a decrease in *spindle frequency* (Figure S6A; interval: $F(2, 44) = 7.89$, $p = .001$, $\eta p^2 = .26$; memory vs. control: $p = .09$; memory vs. rest: $p = .13$; control vs. rest: $p = .006$; visit: $F(1, 22) = .44$, $p = .52$, $\eta p^2 = .02$; visit x interval: $F(2, 44) = 2.03$, $p = .14$, $\eta p^2 = .08$) and an increase in *spindle amplitude* (Figure S6B; interval: ($F(2, 44) = 6.98$, $p = .002$, $\eta p^2 = .24$; visit: $F(1, 22) = .22$, $p = .64$, $\eta p^2 = .01$; visit × interval: $F(2, 44) = 5.89$, $p = .005$, $\eta p^2 = .21$) as compared to both memory cue and no stimulation. The spindle amplitude effect described above was greater for the associated as compared to the unassociated condition (associated visit, interval effect: $F(2, 46) = 8.64$, $p < .001$, $\eta p^2 = .27$; memory vs. control: $p = .04$; memory vs. rest: $p = .17$; control vs. rest: $p = .006$; unassociated visit, interval effect: $F(2, 44) = 3.10$, $p = .06$, $\eta p^2 = .12$). No such effects were observed for *spindle density* (Figure S6C; interval: $F(2, 44) = .28$, $p = .76$, $\eta p^2 = .01$; visit: $F(1, 22) = 1.34$, $p = .26$, $\eta p^2 = .06$; visit x interval: $F(2, 44) = 1.27$, $p = .29$, $\eta p^2 = .05$).

Taken together, these results suggest that auditory stimulation during sleep increased *SO density* irrespective of whether the auditory cue carried or not memory content and whether memories were associated or not during learning. Interestingly, control sounds that did not carry memory information induced an overall decrease in *spindle frequency* and an increase in *spindle amplitude*, the latter effect being greater under high memory load, i.e., during the associated visit when two memory traces are potentially reactivated as compared to one.

### 5. Competition between memory domains for consolidation resources

Our confirmatory analysis revealed an unexpected disruptive effect of direct TMR on the consolidation of the object sequence. One possible explanation put forward in previous research (Frey and Morris 1997) is a competition for consolidation resources. Previous research with the current paradigm in which object sequences were learned after the motor sequences (Temudo et al. 2026) has shown that object learning is disrupted by the previous motor learning task which potentially weakened the encoding of the object memory. One could therefore argue that the stronger motor sequence memories received preferential allocation of consolidation resources relative to the more weakly encoded object memories (Frey and Morris 1997). Under this account, one would expect a trade-off between domains, such that greater TMR index for the motor task is associated with lower TMR index for the object task irrespective of visits (higher order association or not). To test this prediction, we averaged TMR index across visits within each task (motor, object) and examined correlations between motor and object gains. No significant correlation was observed ($r = -.03$, $p = .88$). Together, the present behavioral findings do not support the notion that the observed disruption of the directly reactivated object sequence was driven by competition for resources between motor and object memories (but see main text for brain-behavior exploratory analyses).

### 6. Deviations from pre-registration and justifications

**(1) Auditory-evoked responses were obtained by segmenting the data into epochs time-locked to auditory cue onset (from –1 to 3 s relative to auditory cue onset)**

Epoch segments for the auditory-cue locked analyses were pre-registered as -2.5 to 2s relative to auditory cue onset. We enlarged the post-cue analysis window to accommodate for the longer duration of the auditory cue used in the current research (i.e., 500ms) as compared to our previous work (i.e., 100ms) on which the pre-registration was based. Previous research using similar cue duration reported later responses that could potentially be missed with the pre-registered window (Cairney et al., 2018)

**(2) Individual ERPs and TRFs computed on each channel were baseline corrected by subtracting the mean amplitude from –0.3 to –0.1 s relative to cue onset**

Baseline correction was registered from -2.5 to -2s. We adopted a similar baseline correction as in the study mentioned above using auditory cues of a similar duration as in the present study (Cairney et al., 2018).

### 7. Generation

Analyses of the generation tasks demonstrated that participants successfully reproduced the motor and object sequences with high accuracy. Specifically, the mean percentage of correct ordinal positions, i.e., the percentage of items (movements or objects) generated in the correct temporal position in the sequence and the percentage of correct transitions generated was on average greater than 89% across all sequences, task conditions and visits. A repeated-measures ANOVA revealed no significant differences in the percentage of correct temporal positions or correct transitions between reactivated and non-reactivated sequences (position: $F(1, 23) = .30$, $p = .59$, $\eta_p^2 = .01$; transition: $F(1, 23) = 1.00$, $p = .33$, $\eta_p^2 = .04$) or between task (position: $F(1, 23) = 1.26$, $p = .27$, $\eta_p^2 = .05$; transition: $F(1, 23) = 1.94$, $p = .18$, $\eta_p^2 = .08$) or visits (position: $F(1, 23) = .09$, $p = .76$, $\eta_p^2 = .004$; transition: $F(1, 23) = .33$, $p = .57$, $\eta_p^2 = .01$). There were also no significant interactions between these factors (visit x task; position: $F(1, 23) = .03$, $p = .87$, $\eta_p^2 = .001$; transition: $F(1, 23) = .65$, $p = .43$, $\eta_p^2 = .03$; visit x sequence; position: $F(1, 23) = 1.52$, $p = .23$, $\eta_p^2 = .06$; transition: $F(1, 23) = 1.02$, $p = .32$, $\eta_p^2 = .04$; task x sequence; position: $F(1, 23) = 2.0$, $p = .17$, $\eta_p^2 = .08$; transition: $F(1, 23) = 1.87$, $p = .19$, $\eta_p^2 = .08$; visit x task x sequence; position: $F(1, 23) = .13$, $p = .72$, $\eta_p^2 = .006$; transition: $F(1, 23) = .00$, $p = .97$, $\eta_p^2 = .00$). These results indicate that participants developed explicit knowledge of the series of objects and movements regardless of the reactivation condition, task or whether the sequences were associated or not.

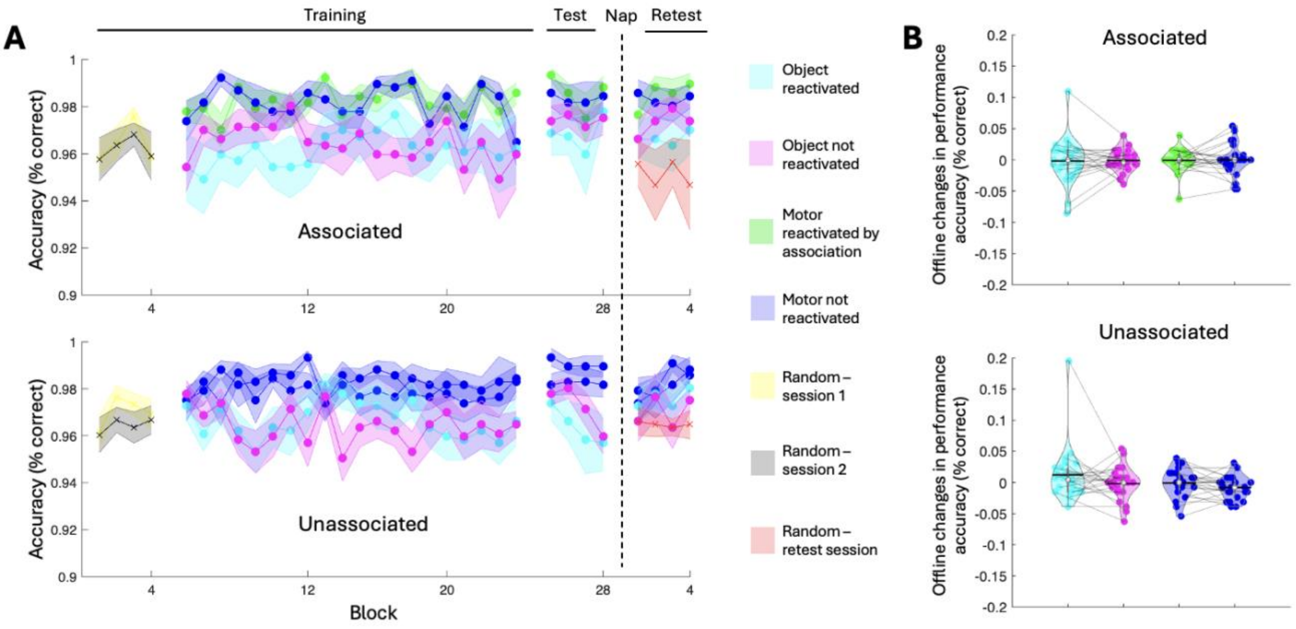


**Figure S1.** *Performance accuracy.* **A.** *Performance accuracy during training* (% correct responses) across blocks for each sequence (reactivated /non reactivated), each task condition (motor/object) and each visit (associated/unassociated). Crosses and circles indicate blocks of the random and sequence conditions, respectively. Shaded error bars=SEM, dotted line indicated when the nap occurred (N = 24). **B.** *offline changes in performance accuracy* (% correct) were calculated as the average accuracy across the four test blocks pre-nap minus the average across the four retest blocks post-nap, for both sequences (reactivated/non-reactivated) of each task condition (motor/object) from both visits (associated/unassociated). Gains were not significantly different between visits, task conditions or sequences. Colored circles represent individual subject data, jittered in arbitrary distances on the x-axis to increase perceptibility. Grey lines map individual subject datapoints across conditions. White circles represent medians, and horizontal bars represent means. The shape of the violin depicts the kernel density estimate of the data.

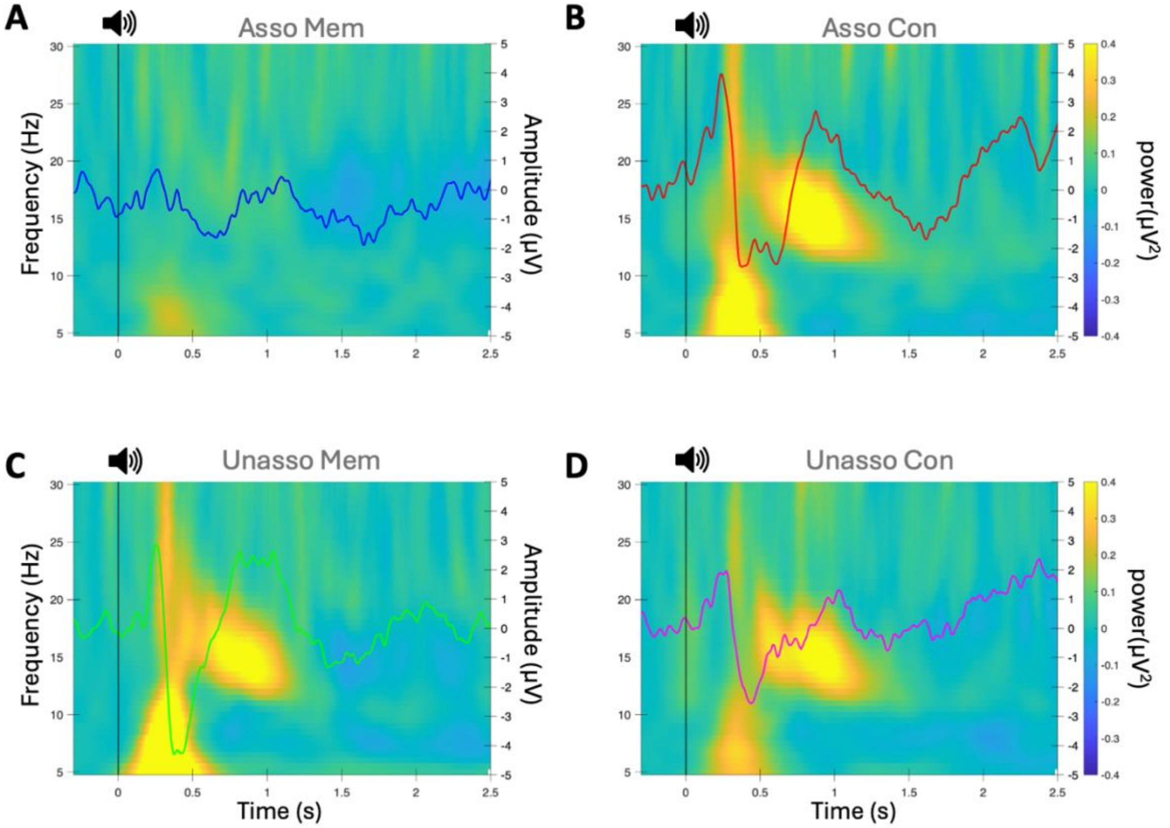


**Figure S2.** *Auditory cue-evoked responses during sleep for each cue and visit condition*. Event-related potentials (ERPs; line plots displayed for channel Cz) are overlaid on time–frequency representations (TFRs; 5–30 Hz) of EEG power (displayed for channel Cz), time-locked to auditory cue onset (0 s; black vertical line). **A.** Evoked responses for the memory (Mem, blue) cue in the associated (Asso) visit**. B.** Evoked responses for the control (Con, blue) cue in the associated visit**. C**. Evoked responses for the memory (green) cue in the unassociated (Unasso) visit**. D.** Evoked responses for the control (magenta) cue in the unassociated visit.

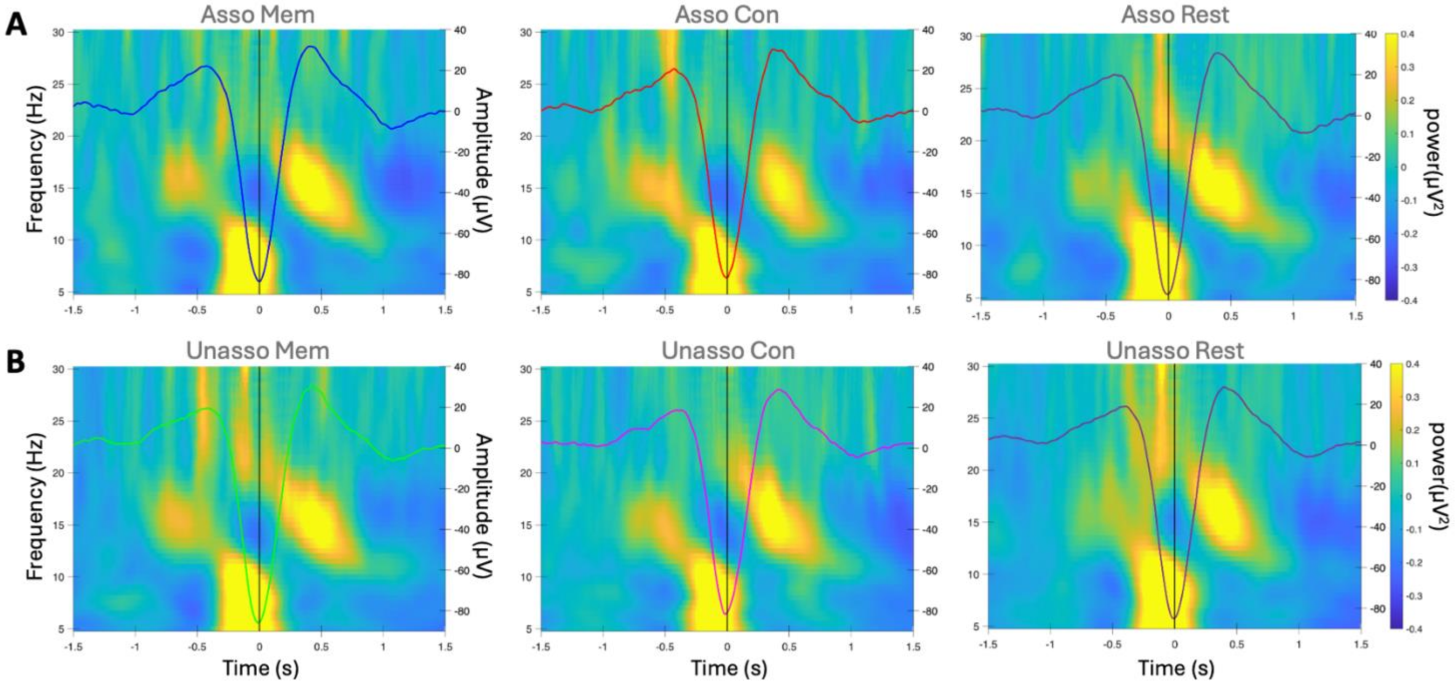


**Figure S3.** *SO-locked responses during sleep for each condition.* Event-related potentials (ERPs; line plots displayed for Fz channel) overlaid on time–frequency representations (TFRs; 5–30 Hz) of EEG power (displayed for channel Fz), time-locked to SO trough (0 s; black vertical line). **A.** Associated visit (Asso, N=24)**;** *left panel*; memory cue (Mem, blue); *middle panel*; control cue (Con, red); *right panel*; rest (purple; no stimulation) **B.** Unassociated visit (Unasso, N=23)**;** *left panel*; memory cue (green); *middle panel*; control cue (magenta); *right panel*; rest (purple; no stimulation).

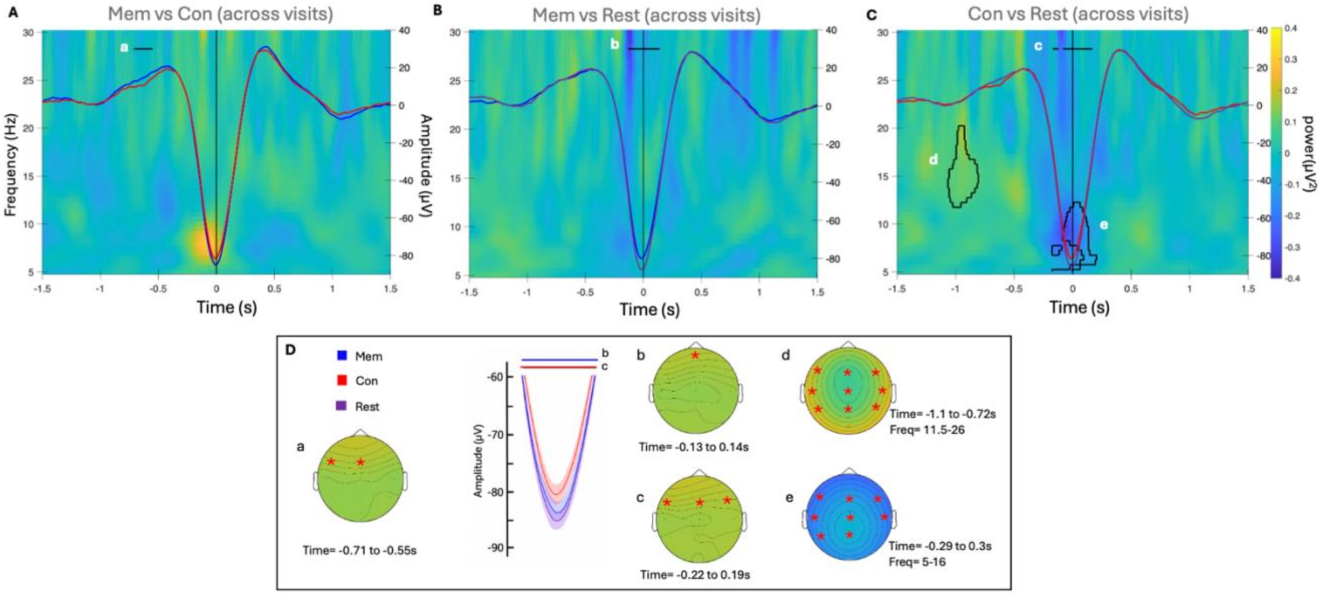


**Figure S4**. *SO-locked responses during sleep across associated and unassociated visits (N=23¥).* Event-related potentials (ERPs; line plots displayed for Fz channel [panel A & C] and FPz [panel B]) overlaid on time–frequency representations (TFRs; 5–30 Hz) of EEG power (displayed for channel Fz [panel A & C] and FPz [panel B]), time-locked to SO trough (0 s; black vertical line). Mem: memory, Con: control. **A.** Difference between memory (blue) and control (red) cue stimulation showing a significant spatiotemporal cluster (a) from −0.71 to -0.55 s (p = .03, Cohen's d = .96; cluster spanning Fz & F3) indicating greater SO amplitude during memory cue compared to control cue stimulation (see panel D for topography of this effect). **B.** Difference between memory cue stimulation and rest (purple) showing a significant spatiotemporal cluster (b) from −0.13 to 0.14 s (p = .01, Cohen's d = 1.08; channel FPz) indicating greater SO amplitude during rest compared to memory cue stimulation (see panel D for topography of this effect and for inset zooming on SO trough). **C.** Difference between control cue stimulation and rest showing significant spatiotemporal cluster (c) from -0.22 to 0.19 s (p= .002, Cohen's d = 1.08; spanning frontal channels) indicating greater SO amplitude for rest compared to control cue stimulation intervals. TF analyses revealed significant spatiotemporal-frequency cluster (d) from −1.1 to −0.72 s, 11.5–26 Hz (p = .004, Cohen's d= 1.05, spanning all channels excluding FPz & Oz) showing greater sigma power 1s before the SO-trough during control stimulation as compared to rest and (e) from −0.29 to 0.3 s, 5–16 Hz (p = .002, Cohen's d= .72; spanning all channels excluding FPz,P4 & Oz) showing greater power during rest compared to control stimulation at the SO-trough (see panel D for topography of this effect and for inset zooming on SO trough). Black horizontal lines indicate time points belonging to significant spatiotemporal clusters in SO amplitude difference identified through cluster-based permutation tests. Black contours in the TFR maps highlight significant spatial time–frequency clusters. **D.** Inset showing zoom on SO trough across visits where the rest condition [purple] showed greater SO amplitude compared to the two auditory stimulation conditions (memory [blue] and control [red]; shaded regions represent SEM). Topographical maps display the spatial distribution of all CBP effects, with red stars marking channels included in each significant cluster**.** ¥ Data excluded for one participant for missing data in the unassociated visit due to experimental error resulting in stimulation triggers not being recorded for this participant.

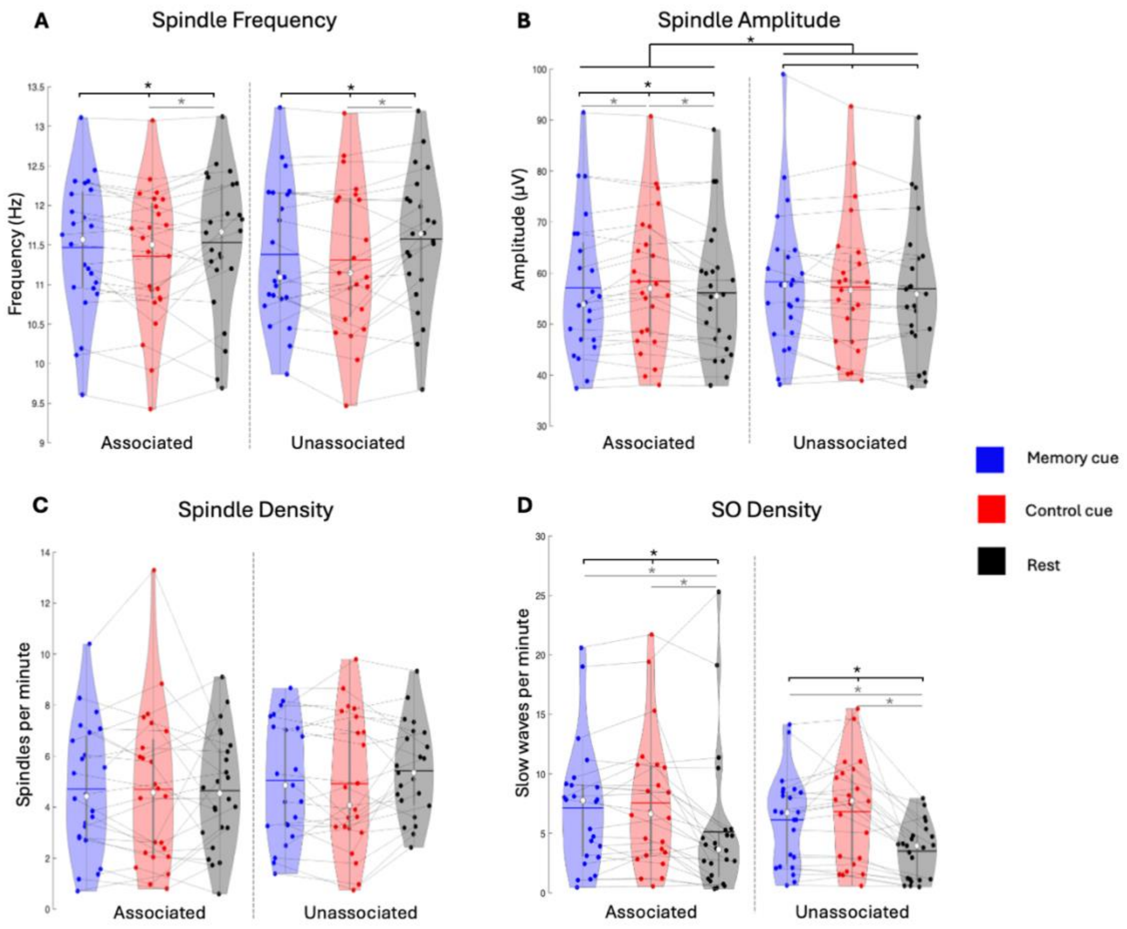


**Figure S5.** *Sleep event characteristics in the different stimulation intervals and visits.* **A.** Spindle frequency (Hz) was lower for control cue stimulation during both visits (interval: $F(2, 44) = 7.89$, $p = .001$, $\eta p^2 = .26$; memory vs. control: $p = .09$; memory vs. rest: $p = .13$; control vs. rest: $p = .006$ across visits). **B.** Spindle amplitude (µV) was greater for control cue stimulation in the associated visit (interval: ($F(2, 44) = 6.98$, $p = .002$, $\eta p^2 = .24$; visit × interval: $F(2, 44) = 5.89$, $p = .005$, $\eta p^2 = .21$; associated visit, interval effect: $F(2, 46) = 8.64$, $p < .001$, $\eta p^2 = .27$; memory vs. control: $p = .04$; memory vs. rest: $p = .17$; control vs. rest: $p = .006$). **C.** Spindle density (number of spindles per minute of stimulation / rest intervals) was not modulated by stimulation interval or visit **D.** Slow oscillation (SO) density (number of slow waves per minute of stimulation / rest intervals) was greater for auditory stimulation intervals compared to rest (interval: $F(2, 44) = 46.27$, $p < .001$, $\eta p^2 = .68$; memory vs. control: $p = .12$; memory vs. rest: $p < .001$; control vs. rest: $p < .001$ across visits). Intervals: memory cue stimulation (blue), control cue stimulation (red) and no stimulation (rest; black) for the associated (N=24) and unassociated visits (N=23[¥]). Colored circles represent individual data, jittered in arbitrary distances on the x-axis to increase perceptibility, white circles represent medians, and horizontal bars represent means. The shape of the violin depicts the kernel density estimate of the data. Black bars with asterisk indicate significant main effect of condition and interactions (across visit), and grey bars with asterisk Indicate significant pairwise differences between intervals ($p < .05$). [¥]Data is missing for one participant in the unassociated visit due to experimental error resulting in stimulation triggers not being recorded for this participant.

| Associated | | |
|---|---|---|
| Set | Motor | Object |
| 1 | seq A: 3,4,1,2,1,3,4,2 | seq A: car, saw, pineapple, giraffe, banana, plane, axe, elephant |
| | seq B: 4,1,2,3,3,2,4,1 | seq B: hammer, pear, camel, bus, helicopter, zebra, screwdriver, orange |
| 2 | seq C: 2,4,1,3,4,3,1,2 | seq C: elephant, axe, banana, plane, saw, car, pineapple, giraffe |
| | seq D: 4,3,2,3,4,1,1,2 | seq D: screwdriver, helicopter, zebra, bus, hammer, orange, pear, camel |
| 3 | seq E: 1,3,1,4,2,3,4,2 | seq E: pineapple, plane, banana, saw, elephant, car, axe , giraffe |
| | seq F: 2,4,4,2,1,3,1,3 | seq F: camel, screwdriver, hammer, zebra, pear, helicopter, orange, bus |
| **Unassociated** | | |
| 1 | seq A: 3,4,1,2,1,3,4,2 | seq C: elephant, axe, banana, plane, saw, car, pineapple, giraffe |
| | seq B: 4,1,2,3,3,2,4,1 | seq D: screwdriver, helicopter, zebra, bus, hammer, orange, pear, camel |
| 2 | seq A: 3,4,1,2,1,3,4,2 | seq E: pineapple, plane, banana, saw, elephant, car, axe , giraffe |
| | seq B: 4,1,2,3,3,2,4,1 | seq F: camel, screwdriver, hammer, zebra, pear, helicopter, orange, bus |
| 3 | seq C: 2,4,1,3,4,3,1,2 | seq A: car, saw, pineapple, giraffe, banana, plane, axe, elephant |
| | seq D: 4,3,2,3,4,1,1,2 | seq B: hammer, pear, camel, bus, helicopter, zebra, screwdriver, orange |
| 4 | seq C: 2,4,1,3,4,3,1,2 | seq E: pineapple, plane, banana, saw, elephant, car, axe , giraffe |
| | seq D: 4,3,2,3,4,1,1,2 | seq F: camel, screwdriver, hammer, zebra, pear, helicopter, orange, bus |
| 5 | seq E: 1,3,1,4,2,3,4,2 | seq A: car, saw, pineapple, giraffe, banana, plane, axe, elephant |
| | seq F: 2,4,4,2,1,3,1,3 | seq B: hammer, pear, camel, bus, helicopter, zebra, screwdriver, orange |
| 6 | seq E: 1,3,1,4,2,3,4,2 | seq C: elephant, axe, banana, plane, saw, car, pineapple, giraffe |
| | seq F: 2,4,4,2,1,3,1,3 | seq D: screwdriver, helicopter, zebra, bus, hammer, orange, pear, camel |

**Table S1**. *Motor and object sequence (seq) sets used in associated and unassociated visits***.**
Each set had two motor sequences (session 1) and two object sequences (session 2). In the Associated visit, each motor sequence was paired with the associated object sequence (e.g., motor seq A paired with object seq A) , such that each finger in the motor sequence was associated to a particular object category in the object sequence. In the Unassociated condition, each motor sequence was combined with an unassociated object sequence (e.g., motor seq A and object seq C), such that each finger in the motor sequence was associated to objects from different categories in the object sequence. Colors mark each sequence (A–F) so that associated sequences can be followed across conditions. Sequence sets were counterbalanced across visits and participants. An example of combination for one individual is set 1 for the associated visit (motor sequences A and B paired with object sequences A and B, respectively) and set 4 for the unassociated visit (motor sequences C and D paired with object sequences E and F, respectively).

| | Associated Visit | Unassociated Visit | Stats |
|---|---|---|---|
| **Time allowed to sleep** | 88.8 min [84.8 – 92.8] | 87.8 min [84.3 – 91.4] | t(23)=0.4, p=0.7 |
| **Total Sleep Time** | 68.7 min [61.0 – 76.3] | 70.1 min [64.7 – 75.6] | t(23)=-0.5, p=0.6 |
| **Sleep Efficiency** | 77.8% [69.3 –86.2] | 80% [74.4 –85.6] | t(23)=-0.8, p=0.5 |
| **NREM1 Latency** | 4.5 min [2.9 – 6.1] | 4.5 min [3.2 – 5.8] | t(23)=0.04, p < 1 |
| **Time awake** | 17 min [11 – 23] | 15.6 min [11.4 – 19.8] | t(23)=0.6, p=0.5 |
| **Time in NREM1 Sleep** | 7.9 min [5.3 – 10.5] | 6.6 min [5.1 – 8.1] | t(23)=1, p=0.3 |
| **Time in NREM2 Sleep** | 33.3 min [26.2 – 40.4] | 36 min [30.8 – 41.2] | t(23)=-1.1, p=0.3 |
| **Time in NREM3 Sleep** | 23.1 min [16 – 30.3] | 23.8 min [18.1 – 29.4] | t(23)=-0.2, p=0.8 |
| **Time in REM Sleep** | 12.2 min [7.5 – 16.9] | 10.4 min [6.5 – 14.2] | t(23)=0.7, p=0.5 |
| **Participants reaching** | | | |
| NREM3 sleep | N = 24 | N = 23 | |
| REM sleep | N = 15 | N = 18 | |

**Table S2***. Sleep characteristics of the experimental nap***.** Values are means [lower and upper limit of the 95% Confidence Interval - CI]. Sleep efficiency was computed as the percentage of time spent asleep (Stage 2 and 3 and REM sleep) relative to the total time in bed (specifically, from lights off to lights on). The stats column shows the results of paired t-tests comparing the data from the associated and unassociated visits. NREM=non-rapid eye movement, REM=rapid eye movement.

| | Associated visit | | | Unassociated visit | | | Stats |
|---|---|---|---|---|---|---|---|
| **Number of Auditory cues** | *All cues* | *Memory cues* | *Control cues* | *All cues* | *Memory cues* | *Control cues* | |
| During all stages | 358.5 [322.1 – 395] | 179.8 [161.9 – 197.6] | 178.8 [159.6 – 198] | 345.4 [293.7 – 397] | 173.1 [146.8 – 199.4] | 172.3 [146.4 – 198.2] | t(22)=-0.4, p=0.7 |
| During wake | 5.1 [1.6 – 8.5] | 2.4 [0.3 – 4.5] | 2.7 [1.1 – 4.3] | 3.9 [0.2 – 8] | 2.1 [1 – 5.2] | 1.8 [0.4 – 3.3] | t(22)=0.6, p=0.6 |
| During NREM1 | 2.7 [0 – 6.4] | 0.8 [0.3 – 1.9] | 1.9 [0.7 – 4.5] | 0.3 [0.1 – 0.6] | 0 [0 – 0] | 0.3 [0.1 – 0.6] | t(22)=1.4, p=0.2 |
| During NREM2-3 | 348.6 [309.6– 387.7] | 176.3 [157.5 – 195.1] | 172.3 [151.5 – 193.1] | 337.9 [287 – 388.9] | 169.8 [143.5 – 196.1] | 168.1 [142.8 – 193.4] | t(22)=-0.6, p=0.6 |
| During REM | 2.1 [0 – 5.2] | 0.3 [0.1 – 0.6] | 1.9 [1.2 – 4.9] | 3.3 [1 – 5.5] | 1.2 [0.2 – 2.1] | 2.1 [0.1 – 4.1] | t(22)=-0.8, p=0.4 |
| Accuracy | 96.8% [94.3 – 99.4] | 97.8% [95.7 –99.8] | 95.9% [92.8 – 99] | 93.8% [85.3 – 102.3] | 94% [85.3 –102.6] | 93.6% [85.1– 102.1] | t(22)=-0.9, p=0.4 |

**Table S3**. *Stimulation characteristics.* Values are means [lower and upper limit of the 95% Confidence Interval - CI]. Accuracy calculated as the percentage of auditory cues sent during Stage 2 and Stage 3 sleep. Stats show results from paired t-tests comparing total number of cues ("All cues") presented in associated and unassociated visits. NREM=non-rapid eye movement REM=rapid eye movement. See Methods section in the main text for comparisons between conditions (memory vs. control cues) and visits (associated [N=24] vs. unassociated [N=23]). Data is missing for one participant in the unassociated visit due to experimental error resulting in stimulation triggers not being recorded for this participant.

| Spindles | | | | | | |
|---|---|---|---|---|---|---|
| | Associated | | | Unassociated | | |
| Channel | memory | control | rest | memory | control | rest |
| FPz | 43.29 [32.02 - 54.56] | 39.92 [28.58 - 51.25] | 90.29 [66.58 - 114.00] | 50.32 [37.68 - 62.96] | 48.23 [35.77 - 60.68] | 124.50 [97.15 - 151.85] |
| Fz | 58.38 [44.86 - 71.89] | 58.08 [42.17 - 74.00] | 100.71 [72.47 - 128.95] | 65.26 [47.43 - 83.10] | 62.04 [44.90 - 79.19] | 128.91 [102.22 - 155.60] |
| Cz | 65.12 [50.97 - 79.28] | 60.75 [45.86 - 75.64] | 110.21 [77.46 - 142.96] | 69.22 [51.55 - 86.88] | 67.26 [49.45 - 85.07] | 140.09 [110.66 - 169.52] |
| Pz | 73.54 [58.37 - 88.72] | 69.29 [53.15 - 85.43] | 133.96 [96.07 - 171.85] | 83.30 [66.11 - 100.50] | 78.52 [60.25 - 96.79] | 175.61 [141.54 - 209.68] |
| Oz | 63.71 [49.77 - 77.65] | 60.75 [46.59 - 74.91] | 135.62 [99.50 - 171.75] | 75.13 [59.20 - 91.06] | 69.35 [53.66 - 85.04] | 171.00 [139.20 - 202.80] |
| F3 | 65.96 [50.11 - 81.81] | 62.50 [45.70 - 79.30] | 104.50 [76.63 - 132.37] | 72.05 [53.19 - 90.91] | 69.14 [51.15 - 87.13] | 139.95 [112.23 - 167.68] |
| F4 | 62.38 [47.22 - 77.53] | 60.12 [43.92 - 76.33] | 106.71 [78.04 - 135.37] | 71.32 [53.57 - 89.07] | 68.18 [50.12 - 86.25] | 137.32 [108.45 - 166.18] |
| C3 | 67.75 [53.18 - 82.32] | 63.62 [48.08 - 79.17] | 120.88 [87.29 - 154.46] | 75.14 [57.85 - 92.43] | 73.50 [56.06 - 90.94] | 157.27 [127.41 - 187.13] |
| C4 | 65.21 [50.90 - 79.51] | 63.58 [47.69 - 79.48] | 120.29 [83.17 - 157.41] | 77.09 [60.67 - 93.51] | 71.23 [53.99 - 88.47] | 165.45 [133.68 - 197.23] |
| P3 | 76.08 [59.62 - 92.55] | 71.71 [55.20 - 88.22] | 139.17 [101.59 - 176.75] | 90.23 [73.42 - 107.04] | 84.27 [66.90 - 101.65] | 190.05 [158.87 - 221.22] |
| P4 | 74.46 [58.42 - 90.50] | 70.17 [53.97 - 86.36] | 141.42 [103.09 - 179.74] | 85.77 [69.06 - 102.49] | 79.32 [61.30 - 97.34] | 186.55 [154.88 - 218.21] |
| Slow ocillations | | | | | | |
| | Associated | | | Unassociated | | |
| Channel | memory | control | rest | memory | control | rest |
| FPz | 139.83 [91.80 - 187.87] | 143.50 [95.57 - 191.43] | 176.12 [82.22 - 270.03] | 124.05 [81.70 - 166.39] | 132.27 [89.93 - 174.61] | 127.14 [77.40 - 176.87] |
| Fz | 152.42 [104.46 - 200.37] | 158.54 [106.81 - 210.27] | 183.92 [94.76 - 273.07] | 135.39 [92.00 - 178.78] | 147.43 [103.24 - 191.63] | 144.17 [90.08 - 198.27] |
| Cz | 129.58 [88.92 - 170.25] | 133.00 [89.47 - 176.53] | 155.83 [81.55 - 230.12] | 111.57 [76.81 - 146.32] | 121.09 [86.60 - 155.57] | 123.00 [80.76 - 165.24] |
| Pz | 112.38 [71.82 - 152.93] | 113.38 [71.91 - 154.84] | 131.33 [61.07 - 201.60] | 96.43 [64.93 - 127.94] | 103.83 [71.64 - 136.02] | 99.04 [62.34 - 135.75] |
| Oz | 64.21 [30.78 - 97.64] | 65.57 [32.65 - 98.48] | 79.54 [25.88 - 133.20] | 52.33 [33.93 - 70.73] | 58.32 [37.49 - 79.14] | 54.18 [28.28 - 80.09] |
| F3 | 132.00 [85.32 - 178.68] | 139.79 [88.40 - 191.18] | 164.71 [77.35 - 252.07] | 116.77 [75.53 - 158.02] | 126.73 [85.14 - 168.31] | 122.64 [73.12 - 172.15] |
| F4 | 144.92 [96.71 - 193.12] | 147.75 [97.11 - 198.39] | 174.29 [82.51 - 266.07] | 125.09 [82.05 - 168.13] | 137.64 [93.44 - 181.83] | 133.86 [81.91 - 185.82] |
| C3 | 100.42 [62.43 - 138.41] | 102.17 [60.82 - 143.51] | 122.33 [53.11 - 191.56] | 82.95 [54.28 - 111.62] | 85.50 [58.34 - 112.66] | 88.73 [54.22 - 123.23] |
| C4 | 104.42 [68.22 - 140.61] | 105.08 [66.69 - 143.48] | 130.38 [58.21 - 202.54] | 89.33 [58.92 - 119.74] | 96.90 [67.22 - 126.59] | 92.29 [61.59 - 122.99] |
| P3 | 92.38 [54.81 - 129.94] | 92.00 [52.34 - 131.66] | 112.54 [45.33 - 179.76] | 68.43 [42.90 - 93.96] | 73.67 [47.38 - 99.95] | 75.19 [42.09 - 108.29] |
| P4 | 90.58 [54.59 - 126.57] | 90.62 [52.81 - 128.44] | 113.67 [45.90 - 181.43] | 72.09 [45.20 - 98.98] | 79.59 [51.60 - 107.59] | 73.73 [44.99 - 102.46] |

**Table S4**. *Spindle and slow oscillation count for each channel, condition and visit.* Values are means [lower and upper limit of the 95% Confidence Interval - CI]. Associated (N=24); unassociated (N=23). Data is missing for one participant in the unassociated visit due to experimental error resulting in stimulation triggers not being recorded for this participant. Slow oscillations events were used for the SO-locked ERP and TFR analyses reported in the main text (and see methods in the main text for a comparison of the number of SOs between interval and visit conditions).

| N = 24 (13 females) | Mean [lower-upper CI] |
|---|---|
| Age (yrs) | 24 ranging from 18 to 35 |
| Edinburgh Handedness (Oldfield, 1971) [a] | 83.5 [77.6–89.5] |
| Epworth Sleepiness Scale (Hoddes et al., 1972) [b] | 6.1 [4.8–7.5] |
| Beck Depression Scale (Beck et al., 1996)[c] | 5.1 [3–7.2] |
| Beck Anxiety Scale (Beck et al., 1988) [d] | 3.2 [1.5–4.7] |
| PSQI (Buysse et al., 1989) [e] | 3.8 [3–4.7] |
| Chronoscore (CRQ) (Horne & Östberg, 1976) [f] | 51.5 [47.6–55.5] |

**Table S5**. *Participant demographics and sleep characteristics***.** Values are means [lower and upper limit of the 95% Confidence Interval - CI]. PSQI = Pittsburgh Sleep Quality Index. CRQ = Circadian Rhythm Questionnaire. [a] -100 = completely left-handed, 100 = completely right-handed; [b] 0 = normal, 24 = severe sleepiness [cut off < 10]; [c] 0 = no depression 63 = severe depression [cut off < 17]; [d] 0 = minimal anxiety, 63 = severe anxiety [cut off < 17]; [e] 0 = good sleep quality, 21 = poor sleep quality [cut off < 8]; [f] 16 = extreme evening type, 86 = extreme morning type [cut off 30 < n < 70].

| | Associated Visit | Unassociated Visit | T-test (asso vs unasso) |
|---|---|---|---|
| **Sleep duration (hours)** | | | |
| Night 1 | 8.3 [7.8–8.7] | 8.5 [8–8.9] | $t(23) = -1.0$ , $p = .33$ |
| Night 2 | 8.5 [8.1–8.9] | 8.6 [8.1–9.1] | $t(23) = -.41$ , $p = .69$ |
| Night 3 | 8.2 [7.8–8.5] | 8.4 [8-8.8] | $t(23) = -.96$ , $p = .35$ |
| Repeated measures Anova | | | |
| Night | $F(2,46) = 1.2$; $p = 0.3$; $\eta p^2 = .05$ | | |
| Visit | $F(1,23) = 1.5$; $p = 0.2$; $\eta p^2 = .06$ | | |
| Night x visit | $F(2,46) = .04$; $p < 1$ ; $\eta p^2 = .002$ | | |
| **St. Mary's questionnaire (a)** | | | |
| Quality | 4.8 [4.3–5.2] | 4.4 [4.3–5] | $t(23) = .49$ , $p = .63$ |
| Duration (hours) | 8.5 [8.2–8.9] | 8.6 [8.2–9] | $t(23) = -.31$ , $p = .76$ |
| **Psychomotor Vigilance Task (ms)** | | | |
| Pre-nap | 334.9ms [315.7–354.1] | 338.7ms [317.7–359.7] | $t(23) = -0.4$ , $p = 0.7$ |
| Post-nap | 298.8ms [257.1–340.4] | 312.5ms [281–344.1] | $t(23) = -0.6$ , $p = 0.6$ |
| Repeated measures Anova | | | |
| Session (pre vs. post) | $F(1,23) = 8.6$; $p = .01$; $\eta p^2 = .27$ * | | |
| Visit (asso vs. Unasso) | $F(1,23) = .42$; $p = .52$; $\eta p^2 = .02$ | | |
| Session x visit | $F(1,23) = .16$; $p = .70$; $\eta p^2 = .007$ | | |
| **Stanford sleepiness score** | | | |
| Pre-nap Session | 2.3 [1.9–2.7] | 2.5 [2–2.9] | $t(23) = -.75$ , $p = .46$ |
| Post-nap Session | 2.4 [2–2.7] | 2.1 [1.7–2.5] | $t(23)= 1.5$ , $p = .16$ |
| Repeated measures Anova | | | |
| Session (pre vs. post) | $F(1,23) = .59$; $p = .45$; $\eta p^2 = .03$ | | |
| Visit (asso vs. Unasso) | $F(1,23) = .04$; $p = .84$; $\eta p^2 = .002$ | | |
| Session x visit | $F(1,23) = 3.2$; $p = .09$; $\eta p^2 = .12$ | | |

**Table S6**. *Participants' nocturnal sleep prior to the experiment and vigilance characteristics***.** Values are means [lower and upper limit of the 95% Confidence Interval - CI]. Sleep duration was computed as the mean across the wrist actigraphy and sleep diary recordings for the three nights before the experimental session. St Mary's questionnaire refers to the night prior to the experimental session; [a] 1 = very unsatisfied, 5 = completely satisfied; Repeated measures ANOVA for sleep duration with night (1-3) and visit (associated vs. unassociated) as within-subject factors. Repeated measures ANOVA for PVT and with session (pre-nap vs. post-nap) and visit (associated vs. unassociated) as within-subject factors. T-test for sleep duration, SMS, PVT & SSS show results from paired t-tests comparing data from associated and unassociated visits. *Performance on PVT task was faster post nap compared to pre-nap.
